# Finding disease modules for cancer and COVID-19 in gene co-expression networks with the Core&Peel method

**DOI:** 10.1101/2020.05.27.118414

**Authors:** M. Lucchetta, M. Pellegrini

## Abstract

Diseases imply dysregulation of cell’s functions at several levels. The study of differentially expressed genes in case-control cohorts of patients is often the first step in understanding the details of the cell’s dysregulation. A further level of analysis is introduced by noticing that genes are organized in functional modules (often called pathways), thus their action and their dysregulation may be better understood by the identification of the modules most affected by the disease (aka disease modules, or active subnetworks). We describe how an algorithm based on the Core&Peel method developed originally for detecting protein complexes in PPI networks, can be adapted to detect disease modules in co-expression networks of genes. We first validate Core&Peel for the easier general task of functional module detection by comparison with 42 methods participating in the Disease Module Identification DREAM challenge of 2019. Next, we use four specific disease test cases (colorectal cancer, prostate cancer, asthma and rheumatoid arthritis), four state-of-the-art algorithms (ModuleDiscoverer, Degas, KeyPathwayMiner and ClustEx), and several pathway databases to validate the proposed algorithm. Core&Peel is the only method able to find significant associations of the predicted disease module with known validated relevant pathways for all four diseases. Moreover for the two cancer data sets, Core&Peel detects further nine relevant pathways enriched in the predicted disease module, not discovered by the other methods used in the comparative analysis. Finally we apply Core&Peel, along with other methods, to explore the transcriptional response of human cells to SARS-CoV-2 infection, at a modular level, aiming at finding supporting evidence for drug repositioning efforts.

## 1 Introduction

In a typical systems biology paradigm, large amount of molecular data collected via high throughput ‘omics’ experiments are stored in curated databases then filtered and reorganized in the form of an interaction network among molecular species (for example, co-expression networks are built via measures of the co-expression of genes under a variety of conditions) [1] [2]. Next, such network is analyzed in order to detect interesting phenomena from a biological point of view, potentially relevant for a phenotype of interest or a specific biological process. Biological networks have been found to have a rich modular structure which mediate biological processes and cellular activities. Thus discovering and validating modules within biological networks has become an activity propaedeutic to the discovery of biological mechanisms. Genes in a functional module should act in an highly correlated (or anti-correlated) way in response to cells conditions, as revealed by approaches integrating different levels of ‘omic’ data [3].

In a second paradigm typical of systems biology, we are not so much interested in finding genes that behave similarly across a variety of cell conditions, but in finding genes (or modules) whose behaviour is different between two clearly stated cell conditions in order to find sub-networks that are activated differently and thus characterize at a functional level the differences between the two conditions (*active sub-network detection problem*) ([4],[5],[6],[7],[8]).

Module and active module identification in biological networks is a key component of a full network analysis aimed at exploring issues related to applications such as drug target discovery [9, 10], cancer subtypes classification [11], finding biomarkers for cancer prognosis [12], [13], detection of histone modifications [14], and many others.

This paper demonstrates the performance of the Core&Peel method for these two key problems in systems biology: (a) network module identification and (b) active sub-networks detection. We show that Core&Peel has a performance that matches or surpasses that of current stat-of-the-art approaches, in several measurements, on benchmark diseases, in particular cancer benchmark data sets. Finally, we apply Core&Peel and other state-of-the-art methods to explore the transcriptional signature of the response of human cells to SARS-CoV-2 infection at a modular level. The combined output of these algorithms uncovers several enriched pathways related to current drug repositioning efforts.

The Core&Peel method has been developed in [15] for predicting protein complexes in large protein-protein interaction networks (PPIN). Core&Peel follows a generalist approach, assuming that modules have the topological properties of quasi-cliques and ego-networks. The method makes very mild assumption on the properties of the input network, and of the predicted modules, thus in principle it can be applied quite directly to other community detection problems with a similar flavour. In practice, the list of 42 methods mentioned in [16] for functional module detection, and the the list of 19 methods mentioned in [15] for protein complex prediction, have very little overlap^1^. Thus the nature of the two problems, functional module identification, and protein complex prediction, is such as to require a fresh look as it is not obvious that any algorithm can perform equally well in either case.

## 2 Methods

### 2.1 DREAM challenge

The “Disease Module Identification DREAM challenge” [16, 17] is an open competition to assess disease-module identification methods across heterogeneous biological networks for homo sapiens. In this section, we summarize the data and the methods that were made available by the challenge. We mention a further recent method developed for the module detection problem which uses data from the challenge by Tripathi et al. [18]. We introduce the strategy used to test for the association of the predicted modules with complex traits and diseases using a collection of Genome-Wide Association Study (GWAS) datasets and a tool for module scoring called PASCAL [19]. Finally, we introduce the Gene Ontology (GO) analysis we performed to evaluate the biological coherence of each predicted module.

#### 2.1.1 Co-expression network

Among the six different biological networks available from the DREAM challenge, we decided to focus our attention on the co-expression network. This network is based on Affimetrix HG-U133 Plus 2 arrays extracted from the Gene Expression Omnibus (GEO) [20]. A gene expression matrix of 22268 genes by 19019 samples was obtained and the pairwise Spearman correlation of genes across samples was computed. The network was created with genes as nodes and the correlation coefficients as edge weights. In order to reduce the noise, we removed all the edges with a weight lower than 0.25, obtaining a network with 12477 nodes and 665187 edges.

#### 2.1.2 DREAM methods

Forty-two methods took part of the DREAM challenge and we compared our results with every algorithm (called DREAM methods in this manuscript). These methods are classified into different categories: (i) kernel clustering, (ii) modularity optimization, (iii) random-walk based, (iv) local methods, (v) ensemble clustering, (vi) hybrid methods and (vii) others. For more details see Table S1 in [16, 17]. The method which performed better than the others using the co-expression network was a hybrid method (coded as H2). This method selected different clustering methods based on the leaderboard performance and structural quality scores. The common characteristic of DREAM methods is to identify non-overlapping modules.

#### 2.1.3 A further method: overlapping community detection

Apart from comparing the Core&Peel results with DREAM ones, we took into account a further method developed by the Raman group [18]. This method, like Core&Peel, is able to detect overlapping modules. Briefly, they used two kinds of seed nodes: HITS and spread hubs, which are based on the degree of a node. Consequently for each seed node, they applied a seed expansion algorithm which uses the Personalized PageRank scores to rank the nodes in the neighborhood of a seed node. The nodes were added to the module one by one based on their ranking and the modularity score was re-calculated after the adding of every node. The set of nodes with the maximum modularity formed a module.

We used their code (seed selection and seed expansion) released on GitHub: https://github.com/RamanLab/DiseaseModuleIdentification. We generated the maximum number of seed nodes in both cases; in particular we had 1423 and 1279 seed nodes for HITS and spread hubs, respectively. In order to compare them with Core&Peel results, we generated modules with pairwise Jaccard coefficient lower than different thresholds (the same stratification used for Core&Peel).

#### 2.1.4 Genome-Wide Association Study (GWAS)

A Genome-Wide Association Study (GWAS) is an approach used in genetics to identify genetic variants associated with risk of disease or a particular trait. In particular, this method searches the genome for small variations, called Single Nucleotide Polymorphisms (SNPs), which consist of a single nucleotide alteration. Generally most SNPs have no effect on health or development. But it has been proven that some of them can be associated to a particular disease or phenotype on the basis of statistical analysis.

The DREAM challenge provides a collection of 180 GWAS datasets from public sources. This collection was split into two sets of 76 and 104 GWASs used for the leaderboard step (parameter optimization) and the final evaluation, respectively. We used both sets separately to test the performance of the methods.

#### 2.1.5 Gene and module scoring using Pascal

PASCAL [19] (PAthway SCoring ALgorithm) is a powerful tool for computing gene and pathway scores from SNP-phenotype association p-values. Basically, Pascal uses analytic and numerical solutions to calculate gene and module scores from the SNP p-values correcting for linkage disequilibrium (LD) correlation structure. PASCAL was used to get a score for each method. This score was defined as the number of modules with significant Pascal p-values in at least one GWAS (if a module is significant for multiple GWAS traits, it was counted once). The Pascal p-values were adjusted to control the FDR via the Benjamini-Hochberg procedure and a 5% FDR cutoff was applied. This methodology is used also in [21].

#### 2.1.6 Gene Ontology (GO) analysis

In order to assess the biological coherence and relevance of each predicted module, we performed the GO analysis. Basically we computed the hypergeometric p-value of the association of each module to each GO class. We used the *hypergeom* method implemented in the *scipy.stats* package of Python. For each module we assigned the GO class of lowest p-value. In order to correct for multiple comparisons, we adjusted the p-values using the *p.adjust* R function with *fdr* method which is the Benjamini-Hochberg FDR estimation method. The GO database was downloaded by ftp://ftp.ebi.ac.uk/pub/databases/GO/goa/HUMAN/goa_human.gaf.gz and the Biological Processes (BP) category was selected. Additionally only the genes in the co-expression network were retained and the genes were filtered to remove the annotation with IEA, ND and NAS evidence codes (corresponding to the “Inferred from electronic annotation”, “No biological data available” and “Non-traceable author statement”, respectively).

### 2.2 Core&Peel

In this section we briefly summarize the main features of the algorithm; for more details refer to [15]. Basically Core&Peel aims at identifying dense sub-graphs from large or medium size networks, that are both ego-networks and quasi-cliques. Briefly it constructs a set of neighbors for each node of the graph based on its core number. Afterwards it applies a peeling step which iteratively removes nodes of minimum degree in the graph. The peeling procedure stops when the number of nodes drops below the minimum size threshold or when the density is above or equal the minimum density threshold. Finally the duplicates were removed and if two subgraphs were too similar according to the Jaccard coefficient, the biggest one was retained.

For both our purposes, we assessed the Core&Peel performance optimizing different control parameters. The subgraph minimum density (d) and the maximum Jaccard coefficient (j) varied from 0.5 to 1.0 and the subgraph minimum size (nl) was set to 10 or 20. Instead the filter strategy (f) was fixed to 1 because it is neither too strict nor loose and the distance from the seed node (r) was set to 1.

#### 2.2.1 CRank

CRank [22] is a general approach for prioritizing network communities. It takes a network and detected communities as its input and produces a ranked list of communities, where the high-ranking ones represent the promising candidates for downstream analyses. It is based on the evaluation of four different structural features of each community: (i) the likelihood of the edges, (ii) internal connectivity, (iii) external connectivity and (iv) relationship with the rest of the network. CRank then applies a rank aggregation method to combine these measures in order to produce the final ranking list of communities.

We applied this ranking method on the communities detected by Core&Peel in order to compare them with the modules detected by the algorithms of the DREAM challenge. This aimed at reducing the effect of the overlap of the Core&Peel communities when compared with DREAM method communities which do not contain any common genes between them.

Basically we selected the first k-communities of Core&Peel with the highest CRank scores, where k is the number of communities detected by the DREAM method we want to compare our result with. In other words, Core&Peel results were compared with every DREAM method, taking the same number of communities (in the most of the cases, Core&Peel detects more communities than DREAM methods).

#### 2.2.2 Selection of active sub-networks

The second purpose of this work is assessing the Core&Peel performance in active sub-network detection. We selected only the Core&Peel modules that include genes which have a significant change of their expression in the patients with disease respect to the healthy samples. More specifically, for each case study we calculated the significance of the overlap between each module and the differentially expressed genes (DEGs) through the hypergeometric test. We took into account only the modules with p-value less than or equal to 0.01. The p-values are computed by the *hypergeom* method implemented in the *scipy.stats* (version 1.1.0) package of Python (version 3.7). Finally all the significant modules are merged in one, taking each gene once. We made this decision to facilitate the comparison of our results with the other methods, which in fact did not take into account modules separately.

### 2.3 Transcriptomic data, pre-processing and differential expression analysis

In this section we summarize the datasets used to identify the active sub-networks from the co-expression network and gene expression data. In total seven gene expression data have been analysed, two are from The Cancer Genome Atlas (TCGA), two are from GEO, and the other three from COVID-19 studies. We also describe the differential expression analysis (DEA) we performed to get the DEGs which were used for the active sub-networks selection (Table S4).

#### 2.3.1 TCGA data

We downloaded and pre-processed the RNA-Seq data from TCGA for prostate and colorectal cancer using the *TCGAbiolinks* R/Bioconductor package [23, 24]. Specifically, we filtered out samples with low tumor purity (<60%) through the *TCGAtumor purity* function [25], using a consensus measurement of tumor purity [26]. We normalized the data for GC-content and library size using the *TCGAanalyze Normalization*. Then we used the *TCGAanlayze Filtering* function to remove the genes with low expression across the samples. Lastly, we performed the DEA with the *edgeR* R package [27]. We compared the tumor with the normal samples and a False Discovery Rate (FDR) cutoff was set to 0.01, along with log fold-change lower threshold *|log*(*FC*)*| ≥* 1.

#### 2.3.2 GEO data

We downloaded the microarray data from GEO for asthma (GSE137268) and rheumatoid arthritis (GSE15573). The normalized data were downloaded using the *GEOquery* R package [28]. We converted the probe names to gene symbols using the metadata. However we needed to apply the *collapseRows* function [29], in order to have one representative gene for each group of probes, since some of these map to the same gene name. In particular, this method chooses the gene with the highest mean across the samples as representative.

Differential expression analyses have been carried out with *limma* [30] setting the FDR threshold to 0.05 and FC lower threshold *log*(*FC*) ≥ 1. We decided to increase the FDR threshold respect to that one chosen for the TCGA data in order to obtain an higher number of DEGs.

#### 2.3.3 COVID-19 data

In order to test the active sub-network detection algorithms in COVID-19 cases, we retrieved the DEGs from two different studies. Xiong et al. [31] carried out transcriptome sequencing of the RNAs isolated from the bronchoalveolar lavage fluid (BALF) and peripheral blood mononuclear cells (PBMC) specimens of COVID-19 patients. In both cases they compared the case with healthy patients and we retrieved the DEGs using the pipeline released by authors. Instead, Blanco-Melo et al. [32] infected the human adenocarcinomic alveolar basal epithelial (A549) cells with SARS-CoV-2 (COVID-cells case) and compare them with controls. The authors released only the statistical parameters from the DEA for each gene. The DEGs were selected choosing the FDR less than 0.05 and FC lower threshold *|log*(*FC*)| ≥ 1.

### 2.4 Validation of the active subnetwork

In this section, we sum up some enrichment analyses conducted to evaluate the power of each method to detect pathways associated to the four diseases studied in this work.

#### 2.4.1 Pathway enrichment analysis : Reactome and DisGeNET

We performed the pathway enrichment analysis based on the Reactome database for each regulatory module obtained for each case study. We employed the *enrichPathway* function of the R package *ReactomePA* [33] to retrieve the enriched pathways with adjusted p-values below the 0.05 cutoff. We calculated the number of disease-associated genes in each enriched pathway using the DisGeNET database [34], one of the largest available collections of genes and variants involved in human diseases. Lastly, we calculated the number of enriched pathways with at least two disease-associated genes.

#### 2.4.2 Enrichment analyses based on different databases

Different databases including specific-disease pathways are available. In order to get a broader investigation of these pathways, we decided to use two R packages *enrichR* [35] and *gprofiler2* [36], which use different databases of interest. In the first case, we conducted the analysis using DISEASES [37], OMIM Disease [38] and WikiPathways 2019 Human [39] databases. In the second case, we took into account the KEGG, GO (BP category) and Human phenotype ontology databases [40]. For each analysis we extracted enriched pathways (with adjusted p-value < 0.05) whose description included specific key-words (example: “prostate”, “carcinoma”, “asthma”, “colon” or “rheumatoid”) and compared their adjusted p-values in −log10-scale.

## 3 Results

### 3.1 Core&Peel: detection of disease modules

Core&Peel can detect modules having different density and grade of separation. We optimized our algorithm varying these two parameters. We noticed that some modules have a size higher than 100 genes and to be consistent with the DREAM challenge, we removed these modules. In fact one of the requirement of the challenge is predicting modules no bigger than 100. Figure S1 and Figure S2 represent the number of modules detected by Core&Peel with or without modules larger than 100 genes, respectively.

To assess the Core&Peel performance in detecting disease-modules, we applied the PASCAL algorithm using the leaderboard and final GWAS datasets. Actually we used the leaderboard step to examine which configuration gave the best result. We decided to focus on Core&Peel-r1-nl20-d0.7-f1-j0.8 (Figure 1), since it detects a large number of enriched modules in the leaderboard dataset. Generally when the Jaccard index is from 0.8 and 1.0 there is not a substantial variation in the number of disease modules identified, so we decided to choose the smallest Jaccard index into this range. However when the density is too high (especially equal to 1) very few modules were identified. This is more evident in the leaderboard step, in fact when the final GWAS datasets were used a remarkable improvement can be noticed, probably due to the number of datasets larger than those in the leaderboard.

**Figure 1:**
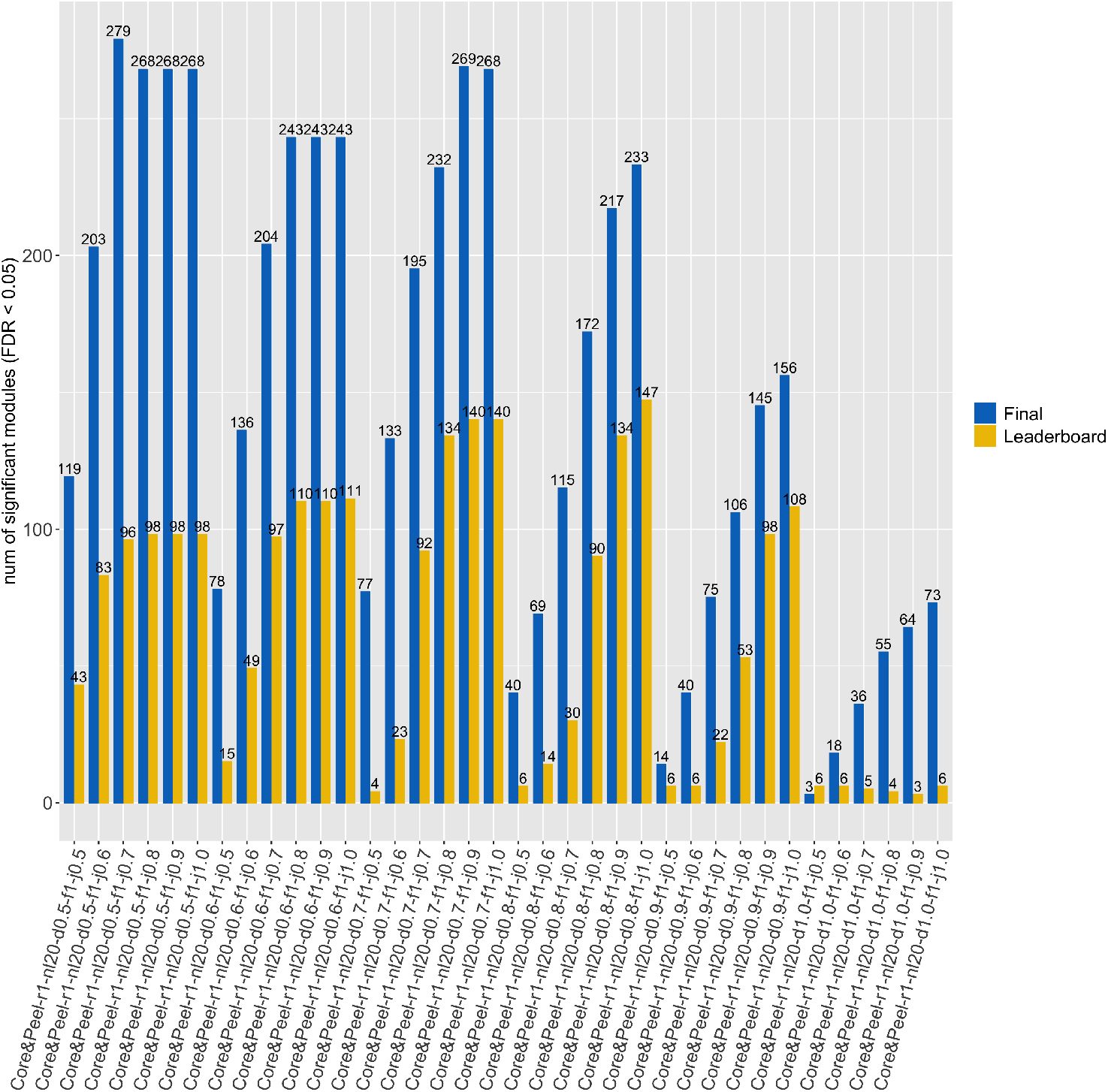
Number of disease-enriched modules detected by Core&Peel using the Leaderboard and Final GWAS datasets. In abscissa the tested configurations of parameters for Core&Peel.

We also noticed that removing the modules with size higher than 100 slightly improves the performance, in fact an equal or larger number of disease modules were detected (compare Figure 1 with Figure S5).

#### 3.1.1 Comparison of Core&Peel with DREAM methods

In order to compare Core&Peel with DREAM methods, we applied the CRank algorithm to the configuration chosen (Core&Peel-r1-nl20-d0.7-f1-j0.8) and selected the highest score communities to obtain the same number of modules of DREAM methods (Figure S3).

The results of leaderboard and final tests for Core&Peel versus the DREAM methods are represented in Figure 2. In most of the cases, Core&Peel can identify more enriched modules. In few cases Core&Peel finds fewer GWAS enriched modules, for example when compared with K5 (Figure 2b) probably due to the low number of predicted modules (equal to 16). Obviously Core&Peel has the advantage to have overlapping modules and this helps to obtain more disease enriched modules for equal number of predicted modules with DREAM methods. This highlights the importance of taking into account modules with genes in common since they are more realistic from a biological point of view, as demonstrated also in [18].

**Figure 2:**
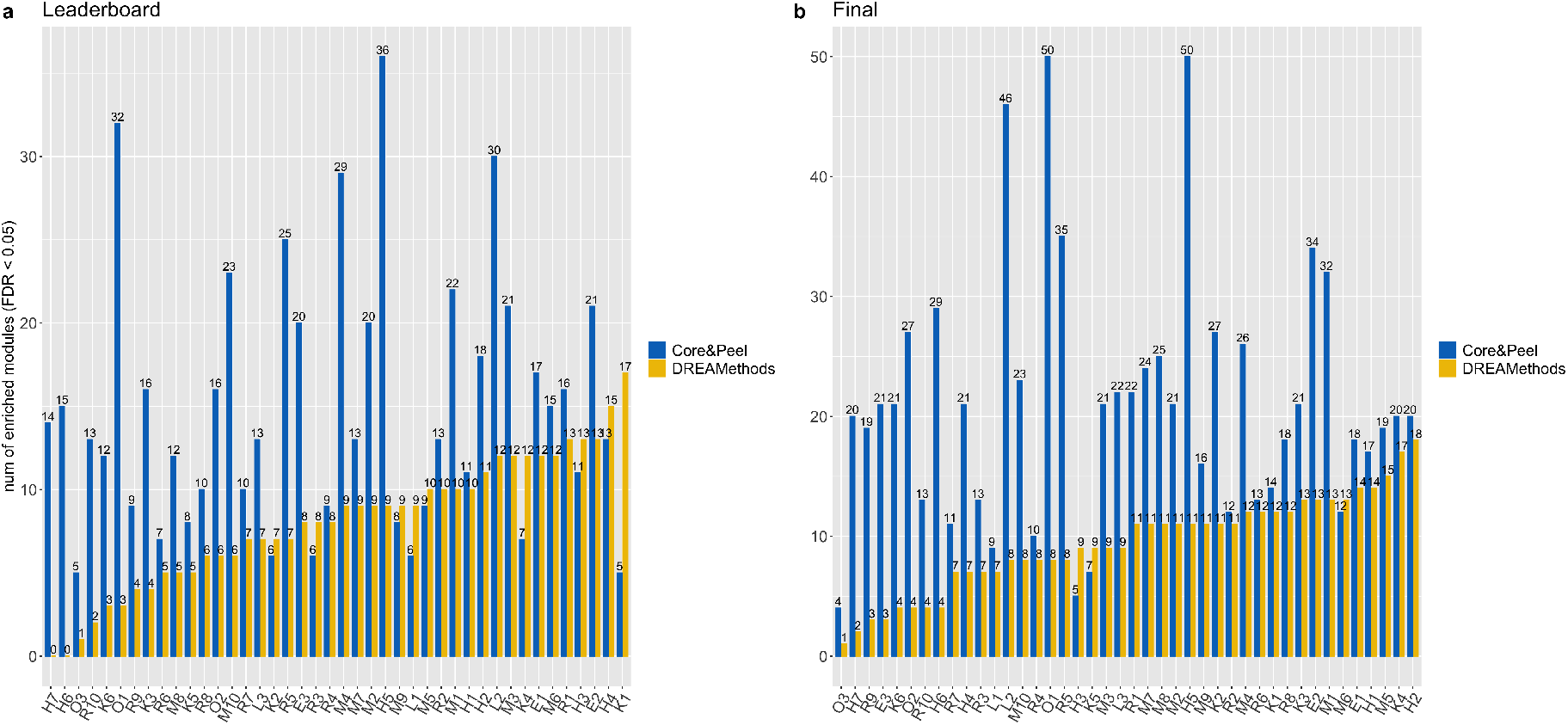
Results of PASCAL tool using leaderboard (a) and (b) final GWAS datasets. Comparison between Core&Peel-r1-nl20-d0.7-f1-j0.8 and all the DREAM methods. The x-axis is in ascending order respect to the number of enriched modules detected by DREAM methods.

Moreover, we performed the GO analysis to evaluate the biological coherence of each predicted module. As before, we applied the CRank ranking to Core&Peel before conducting the enrichment analysis. We compared the number of predicted modules with different thresholds of significance (Table S1) and Core&Peel enables to detect on average more functional modules than DREAM methods; in particular when the FDR threshold is very low.

#### 3.1.2 Comparison between Core&Peel and the overlapping community detection method

For the sake of completeness, we compared Core&Peel with the method [18] described in the Section 2.1.3 which detects overlapping modules like Core&Peel. Also in this case, Core&Peel identifies many more disease-enriched modules (Figure 3). We also noticed that Core&Peel predicts in total much more modules than this method (Figure S4). Consequently we showed the fraction of predicted modules that are enriched (Table 1) and also in this case Core&Peel performs better ^2^. In addition, we analyzed the biological relevance of the predicted modules with GO analysis and Core&Peel achieves the best result (Table S2 and S3).

**Figure 3:**
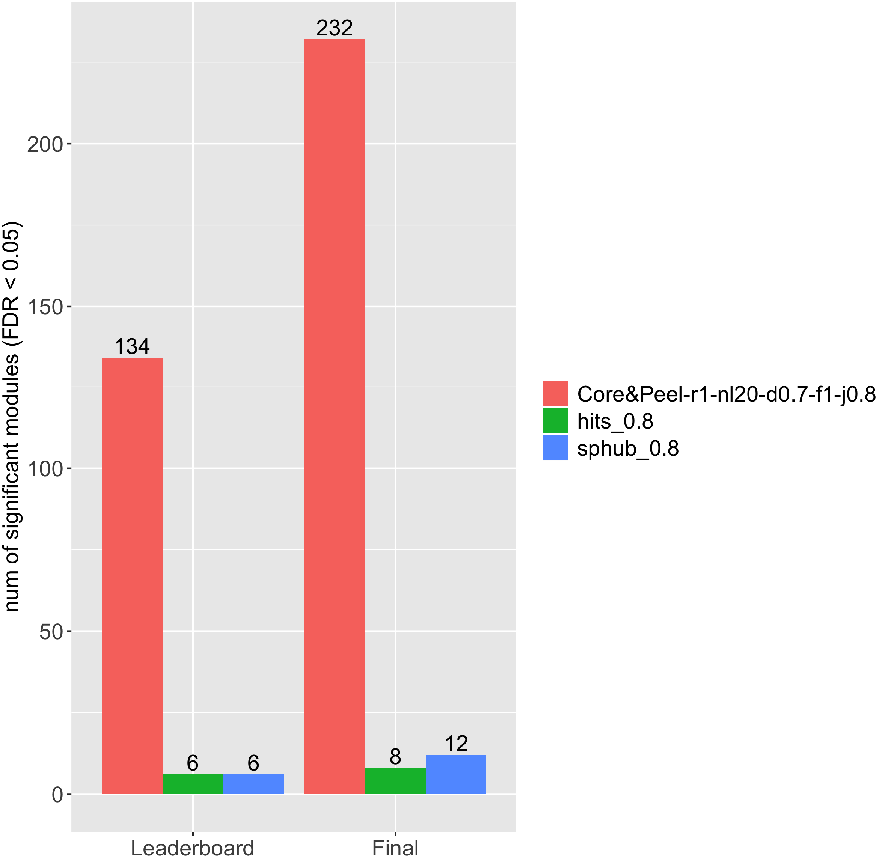
Number of disease enriched modules using both Leaderboard and Final GWAS datasets and comparing Core&Peel-r1-nl20-d0.7-f1-j0.8 with the two options of the method explained in Section 2.1.3. In both cases a Jaccard maximum index of 0.8 was selected to compare them with Core&Peel.

**Table 1:** Fraction of predicted modules that are enriched in the Leaderboard and Final GWAS datasets. The numbers in bold highlight the best approach.

|  | Core&Peel-r1-nl20-d0.7-f1-j0.8 | hits_j0.8 | spread_hubs_j0.8 |
| --- | --- | --- | --- |
| Leaderboard | **0.021** | 0.004 | 0.005 |
| Final | **0.037** | 0.006 | 0.01 |

### 3.2 Core&Peel: detection of active sub-networks

After running Core&Peel, only significant modules have been selected (Section 2.2.2) for each case study. We ran Core&Peel using both 10 and 20 as the subgraph minimum size. We noticed that, in the active sub-network detection problem, Core&Peel with nl = 10 performs better. So we chose the Core&Peel-r1-nl10-d0.7-f1-j0.8 since it is one of the best configurations (same density and Jaccard coefficient) on the DREAM challenge data. Actually this configuration works better in prostate and colorectal cancer cases where the number of DEGs is pretty high, instead when the number of DEGs is low (Table S4), Core&Peel with a smaller density obtained better results. So we decided to use Core&Peel-r1-nl10-d0.5-f1-j0.8 in the asthma and rheumatoid arthritis cases. In COVID-19 case studies, we took into account both Core&Peel configurations.

Core&Peel detected 1270, 141, 49 and 1495 modules in the prostate cancer, asthma, rheumatoid arthritis and colorectal cancer datasets, respectively. As expected, the more are the DEGs (Table S4), the more modules are identified. Since the significant modules were merged in one active subnetwork, we checked if this subnetwork is still significantly enriched with the DEGs; all active subnetworks produced got a p-value < 10^−18^ (data not shown).

#### 3.2.1 Comparison with other active sub-network detection methods

We compared the Core&Peel modules with those detected by the four existing active sub-network identification methods, namely ModuleDiscoverer (MD) [41], KeyPathwayMiner (KPM) [42], ClustEx [43] and Degas [44] (see description on Supplementary Notes Section 3.1). The number of genes in each active subnetwork is reported in Table 2. Generally, regulatory modules of asthma and rheumatoid arthritis include a smaller number of genes than the prostate and colorectal cancer modules; this could be due to the lower number of DEGs. Degas is the method that produced the smallest modules, Core&Peel, ModuleDiscoverer and ClustEx produced the biggest ones, instead.

**Table 2:**
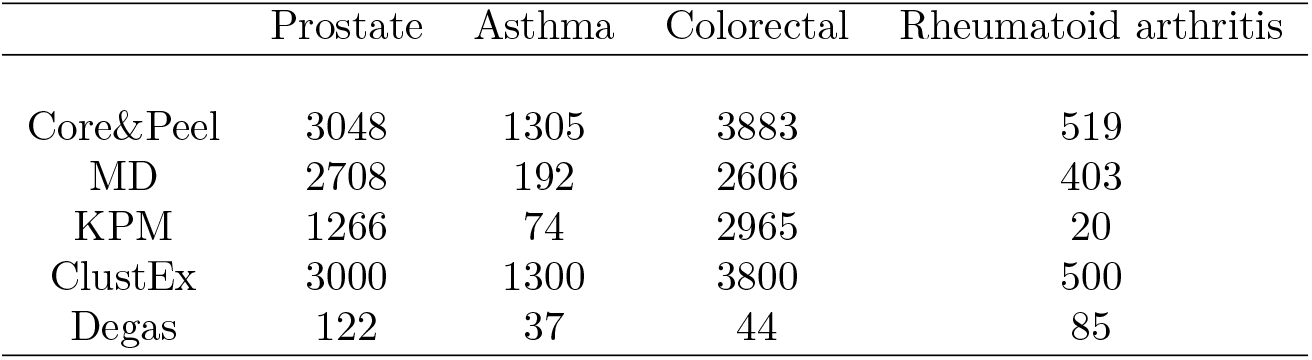
Number of genes in each active sub-network detected by all the methods for the four case studies.

Next, we were interested to compare the modules detected by the different methods from a biological point of view. We compared the enriched pathways (Reactome-based) and plotted the overlap among all the method combinations (Figure 4). We also conducted as baseline the pathway enrichment analysis using the DEGs. Globally, MD, ClustEx and Core&Peel detected a higher number of pathways than Degas, KPM and DEGs. Degas is the method which identified the lowest number of pathways. In particular, in prostate and colorectal cancer cases (Figure 4a and 4b), it does not have any overlap with the other methods. In both cancer cases, Core&Peel and MD share a great part of their pathways (around 40) and a large number of pathways was identified only by ClustEx and MD in prostate (Figure 4a) and colorectal cancer (Figure 4b), respectively. In rheumatoid arthritis case, more than half of the pathways were shared by Core&Peel, ClustEx and MD (Figure 4d) and there is a small overlap for the other combinations. Overall a smaller number of enriched pathways were detected in asthma (see horizontal bars on Figure 4c) than the other three case studies (horizontal bars on Figure 4a, b and d) and we can also notice there are fewer overlapping modules among the methods (Figure 4c). In fact almost the entire set of enriched pathways of ClustEx is not in common with those produced by any other method. Similarly, more than half of the pathway identified by Core&Peel were not detected by the other methods (Figure 4c).

**Figure 4:**
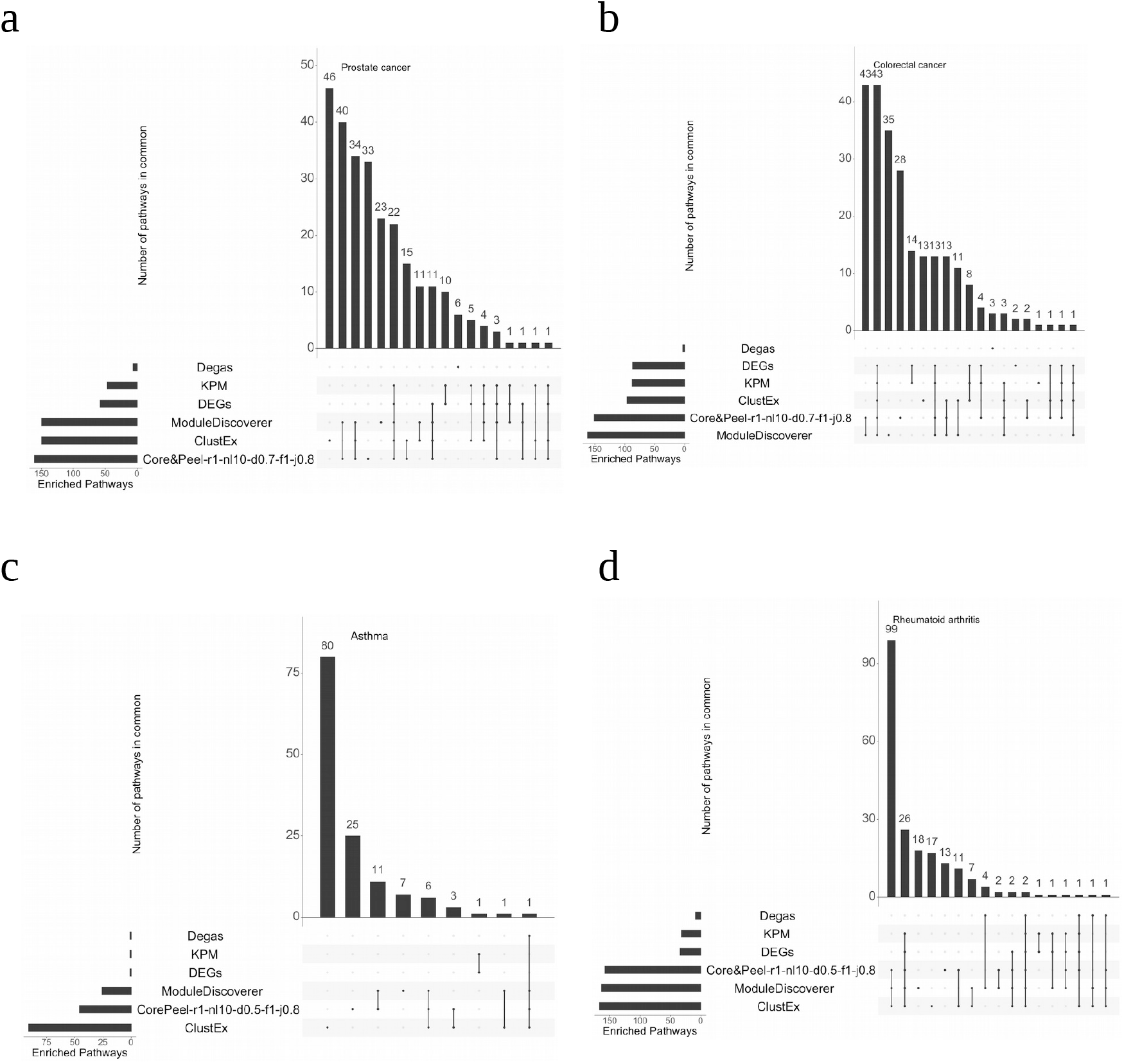
Overlap of enriched pathways among all method combinations in four case studies. The bars on the bottom-left represent the number of enriched pathways detected by the methods. The bars on the main plot represent the number of enriched pathways in common between methods marked with black points on the panel below. The plot was generated by the R package *UpSetR*.

#### 3.2.2 Detection of disease-associated pathways

In order to assess the performance in detecting pathways with genes associated to the disease, we calculated the number of enriched pathways with at least two genes associated to the specific disease (annotated on DisGeNET database). The results are showed in Figure 5. Generally Core&Peel is able to identify a substantial number of pathways. It shows to be a competitive method and it outperforms most of the algorithms. In particular it performs extremely well in asthma case, in which most of the competitors can not detect any pathway with disease-associated genes. In fact Core&Peel and ClustEx are the only methods in able to identify some of them, with Core&Peel detecting a larger number of pathways (Figure 5b). Overall a number more consistent of pathways with disease-associated genes was identified in the two cancer cases (Figure 5a) than asthma and rheumatoid arthritis (Figure 5b). Generally Degas is the method that performs worse in all the case studies (almost zero pathways with disease-associated genes in all cases). Core&Peel, along with ClustEx and MD, is able to identify more pathways with disease-associated genes than DEGs; this highlights how the combination of network analysis and gene expression data can increase the power to detect pathways associated to the disease, which otherwise would not have been identified.

**Figure 5:**
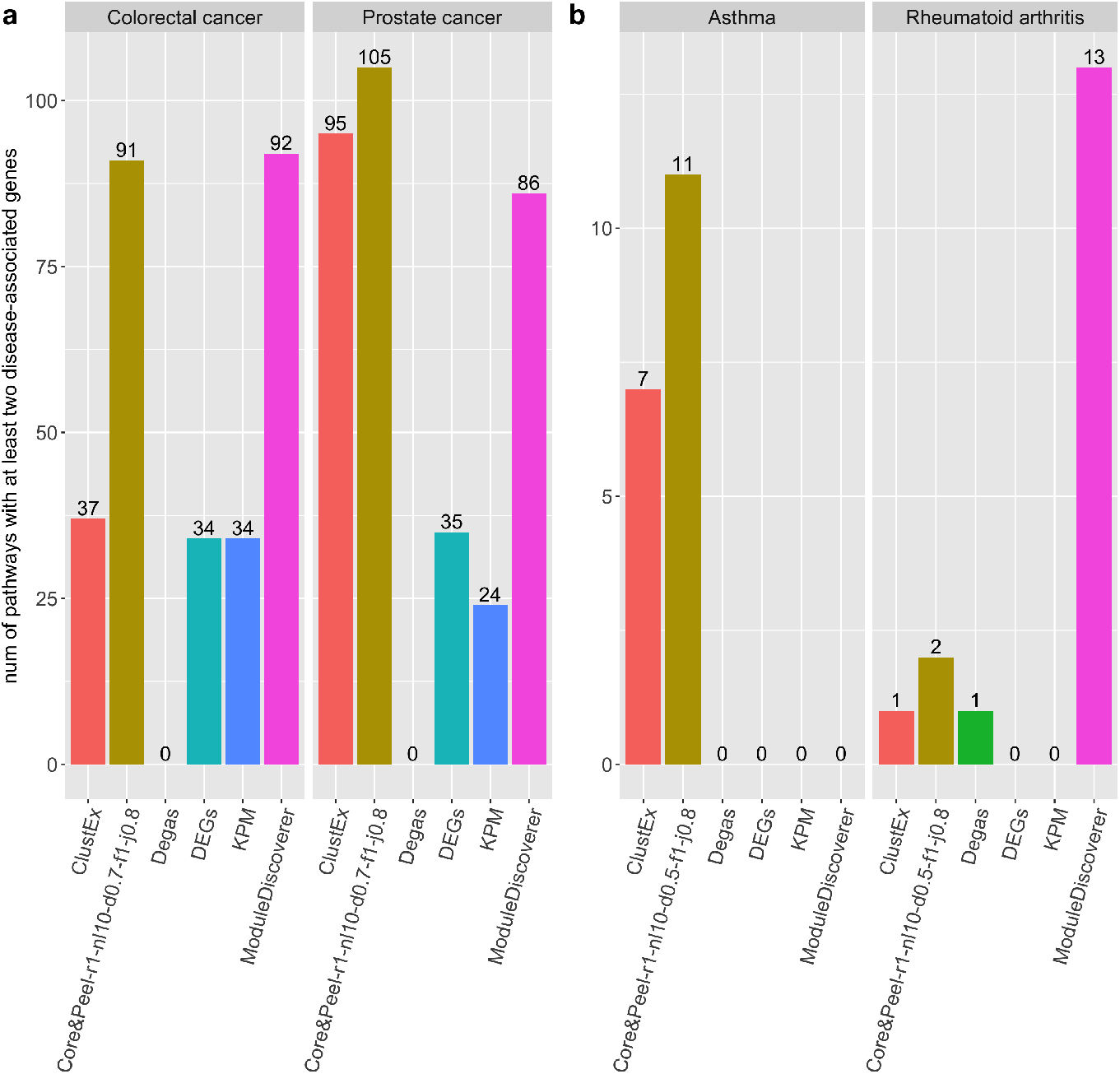
Number of enriched pathways with at least two disease-associated genes. The pathway enrichment analysis was performed using the Reactome database and the disease-gene association was exctracted by the DisGeNET database. (a) Prostate and Colorectal cancer cases; Core&Peel-r1-nl10-d0.7-f1-j0.8 was used. (b) Asthma and rheumatoid arthritis cases where Core&Peel-r1-nl10-d0.5-f1-j0.8 configuration was used.

Finally, we conducted further enrichment analyses using different databases in order to investigate the enrichment with specific-disease pathways. The results are showed in Figure 6. Modules identified by Core&Peel, KPM, ClustEx and MD (along with DEGs) are significantly enriched for the *Carcinoma* pathway in both cancer cases (Figure 6a), suggesting their relevance to cancer in general. More specifically, the Core&Peel module is the only one to be significantly enriched for *prostate cancer* specific pathway (Figure 6d). Moreover, also pathways involved in the prostate morphogenesis (*prostate gland morphogenesis, prostate gland epithelium morphogenesis* and *branching involved in prostate gland morphogenesis*) were significantly enriched and each of them contains some prostate cancer associated genes, suggesting a possible involvement of these pathways in the prostate cancer (Figure 6d). The *epithelial to mesenchymal transition in colorectal cancer* pathway was significantly enriched in Core&Peel module (Figure 6c), giving a colorectal cancer specificity to the Core&Peel module. Moreover, the Core&Peel showed an enrichment for the asthma pathway even if slightly less significant than ClustEx (Figure 6b). No significantly enriched pathways were found for the rheumatoid arthritis case. Finally in Supplementary notes (Section 3.2.2) we reported the same analysis using Core&Peel-r1-nl20-d0.8-f1-j0.8. Also in this case, Core&Peel is able to detect modules enriched for specific prostate and colorectal cancer pathways (*neoplasm of the colon, abnormal prostate morphology, prostate cancer* and *prostate neoplasm*).

**Figure 6:**
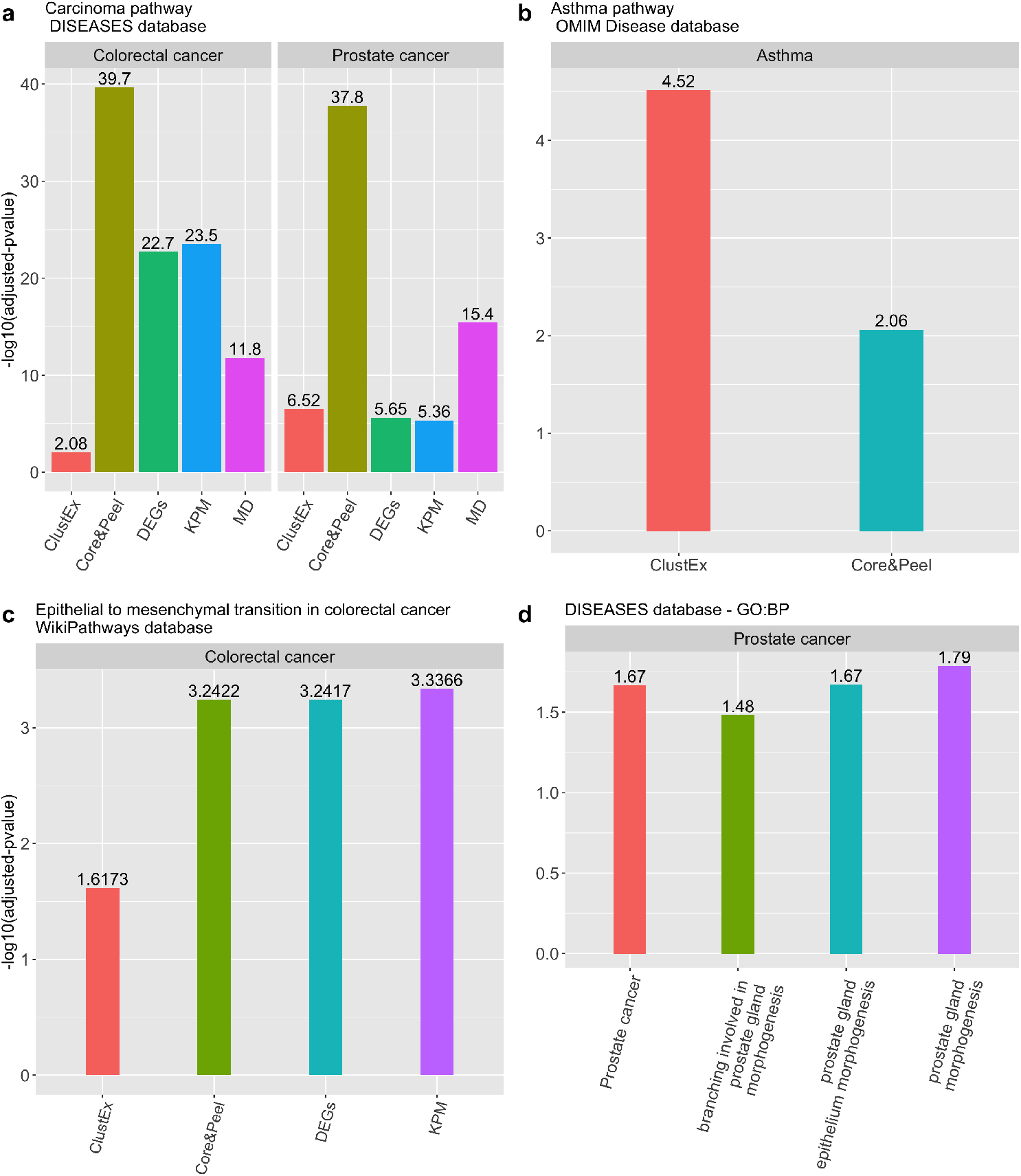
Results of *gProfileR* and *enrichR* analyses. The bars represent the adjusted p-value in log10-scale. (a) The adjusted p-values of Carcinoma pathway annotated in DISEASES database in colorectal and prostate cancer. Only the methods which reached a significant p-value (< 0.05) are showed. (b) Adjusted p-values of asthma pathway annotated on OMIM Disease database. (c) Adjusted p-values of epithelial to mesenchymal transition in colorectal cancer pathway annotated on WikiPathways database. (d) Adjusted p-values of Prostate cancer pathway (annotated on DISEASES database) and prostate gland morphogenesis, prostate gland epithelium morphogenesis and branching involved in prostate gland morphogenesis (annotated on GO).

Overall this analysis has highlighted how Core&Peel is able to find modules specific to the disease.

#### 3.2.3 Comparison with other active sub-network detection methods on COVID-19 data

The numbers of genes in each module detected by the all methods are reported in Table S5. Both Core&Peel configurations (Core&Peel-r1-nl10-d0.5-f1-j0.8 and Core&Peel-r1-nl10-d0.7-f1-j0.8) have been tested in order to have a broader exploration of the enriched pathways. Consequently, two versions of ClustEx have been generated in order to get modules with size comparable with Core&Peel. In the three COVID-19 cases, the biological similarities between methods are different case by case. In BALF case (Figure S6), Degas has detected a considerable number of pathways more than the other methods, and most of them are not in common among othr methods. Core&Peel and ClustEx share large part of the pathways. In PBMC case (Figure S7), many pathways were detected only by ClustEx and a large part of pathways are in common among Core&Peel, MD and Degas. Finally, in COVID-invected cells case (Figure S8), Core&Peel detected the greatest part of pathways and just a few are in common with the others methods.

## 4 Discussion

In this work we explore the potential value of the Core&Peel method for solving two related problems. The first problem is functional module detection within a biological network of interacting genes. The second is the problem of active functional module detection, that is, finding modules within a biological network of interacting genes, that are most affected in a specific disease vs. a normal background state. We initially discuss the two problems separately as the validation methodology is different in either case, however the two aspects of the issue often interact with each other.

### 4.1 Choice of the type of input network

All methods mentioned in this paper require in input a biological network in which nodes represent genes, and edges represent interactions between pairs of genes. Many methods for active sub-network detection have been developed and tested using as input PPI networks (protein-protein interaction networks) [5]. Instead for this study we chose as target (large) gene co-expression network (co-expr) developed within the DREAM challenge project.

There are advantages and supporting evidence that using a co-expr network is a sensible choice. (a) an edge in co-expression network may be due both to direct protein-protein interactions of the proteins associated to the genes, as measured in large scale proteomic essays, but may be also due to indirect co-regulation in which the two genes show correlation because their expression is modulated by a common Transcription Factor or a ncRNA (miRNA, lncRNA, etc..). Thus a richer array of phenomena is captured in a co-expr network. (b) Co-expr networks tend to have more edges and thus to be denser than most PPIN. This is an advantage for density based-methods, as modules tend to be more compact (in other words the average path length within a module is shorter for the same module detected in a denser network). (c) Co-expr networks are usually based on correlation between expression vectors of pairs of genes, across thousands of measurements of gene panel (via microarray or RNAseq technologies). This set up integrates naturally data from different experiments, and different platforms, via standard normalization. The resulting correlation values depend very little on outlier conditions, or variations across experimental conditions, thus represent a robust datum. In contrast, integration of PPI data is somewhat more affected by outlier experiments, and eterogeneous experimental conditions. (d) Huang et al. [45] compare the performance of 21 networks in uncovering pathways associated with a particular disease phenotype, by using 50 literature-based validated pathways. Huang et al. note that larger network size gives higher absolute performance score. Moreover, among seven networks integrating co-expression data, six of them rank in the top seven positions by absolute performance score. (e) The findings in [45] are also consistent with the observations in [16] as the largest average number of disease-related modules is detected (averaging among methods) in the co-expression network (of 1M edges). Slightly less performant is a PPI network which however has more than twice as many edges (about 2.2 M).

On balance, although there is evidence pointing at co-expression networks as an input of choice, both [16] and [45] indicate that usually different types of networks reveal different aspects of the active subnetworks, thus a multi-facet approach should always be considered for specific investigations.

### 4.2 Functional module detection

We compare Core&Peel with about 42 methods that participated in the DREAM challenge project, as well as some recent methods of Tripathi et al. 2019, which allows substantial module overlap. Module overlap is an issue that arises often in the context of network partitions and modularization (as also in graph based community detection). The DREAM challenge did require the competing methods to report non-overlapping modules. This choice has a first effect, since there is no need in the DREAM challenge of a requirement on the maximum number of reported modules. The number of nodes in a network is a natural upper bound on the number of reported modules. It is also somewhat easier to evaluate the enrichment of disjoint modules since one does not have to cope with inflated results, where possibly many high quality overlapping modules differ by just one or two genes, collectively representing quasi-duplicates of the same biological module.

On the other hand, there is ample evidence [46, 47, 48, 49] (see also the the discussion in the DREAM paper [16]) that functional biological modules often share sub-modules and that this sharing has precise implications for the biological interpretation of the processes under study.

When we compare Core&Peel with the output of the DREAM challenge we mitigate the potential bias in the measurements due to overlapping modules by imposing a maximum value of the Jaccard coefficient between any pair of modules, thus avoiding to report quasi-duplicates. Moreover, for the sake of comparison, we rank the modules by intrinsic topological quality (see [22]) and we limit Core&Peel to the top k modules in this ranking when we compare it to a method reporting k modules (for the same input graph).

Within these caveats, we follow strictly the performance measurement methodology of DREAM. We use separately the GWAS leaderboard and the final datasets (see Figure 1). The results on the GWAS leaderboard are used to optimize the choice of the parameters of the Core&Peel method. Figure 1 reports good qualitative concordance of the relative performance of Core&Peel between leaderboard and final GWAS datasets, for a wide range of parameters. There is a discrepancy for the density value *d* = 1.0, which corresponds to detecting full cliques in the input graph. Since even the best co-expr networks are approximations to the true network of interactions, missing edges (false negatives) are to be expected in the input, thus in effect implying that quasi-cliques may be more relevant to functional module detection than full-cliques.

Figure 2 reports the comparison of the absolute number of enriched modules found by Core&Peel (with the selected parameters) versus the DREAM methods. Both for the leaderboard and the final GWAS data, Core&Peel restricted to the top k modules finds more enriched modules most of the times (for final GWAS 39/42 times, for leaderboard GWAS 35/42 times).

Comparing the method of Tripathi et al. 2019 [18], results in Table 1 and Figure 3 show that Core&Peel is able to detect many more significant modules (with a common Jaccard coefficient maximum threshold of 0.8), and that Core&Peel has a larger fraction of the reported modules enriched for GWAS data.

We also performed a second analysis in which we count the number of modules taht are enriched within one (or more) Gene Ontology annotation at a FDR below a fixed threshold (see Tables in suppl. materials S1, S2, and S3). Table S1 reports the number of enriched modules for the DREAM methods, as well as the average number over all DREAM methods. The average number of GO:BP enriched modules for the DREAM methods is comparable to those found by Core&Peel for thresholds 10^−2^, and 10^−3^, however for thresholds form 10^−4^, down to and 10^−7^ Core&Peel has a substantial higher number of enriched modules. At threshold 10^−7^ Core&Peel reports more enriched modules than 39/42 of the DREAM methods.

Comparing the method of Tripathi et al 2019 [18], results in Table S2 and S3 show superior performance of Core&Peel, both in terms of enriched modules for GO categories, and in terms of fraction of enriched modules over the total number of returned modules.

Both the validation with GWAS data and that with GO data give the same picture, showing that Core&Peel is competitive with the best methods in the field on many measured quality functions, even when discounting for the difference in the number of reported modules. Moreover as note in [17] (page 8) the validation via GWAS enrichment uses data that is derived from association studies that are not normally used to define pathways and functional categories found in databases.

### 4.3 Active module detection

Table S2 and S3 show that at threshold 10^−4^ Core&Peel reports 6285 modules, and more than 53% of these (3343) are enriched with one or more GO annotations. This is a very rich structure and we can exploit this richness to attack the problem of active module detection. Similarly to other methods, like ModuleDiscoverer, we use the list of gene differentially expressed (DEG) in case control experiments to compute the enrichment of each module in DEG, and we report as active modules those with an enrichment below a FDR threshold (typically 0.01). Other methods like Degas [44] (which is available the suite Matisse [50]) take a different approach, as the original input graph is annotated with DEGs, and the subnetwork induced by the DEG nodes is used as a seed to discover a minimally connected subgraph that joins all DEGs.

We compare the performance of Core&Peel with 4 algorithms and a baseline (see Table 2) over four test cases (basic statistics in Tables S4 and S5). We have two cancer datasets (prostate and colorectal), one for asthma, and one for rheumatoid arthritis. The number of DEGs in each case varies from a few thousands (for cancer), to a few hundreds (asthma) to a few dozens (rheumatoid arthritis). The performance of Core&Peel is sensitive to the number of DEGs and we get qualitatively different results that may depend mainly on the number of DEGs.

The main validation methodology for the active sub-network detection methods involves finding validated disease-related pathways and testing the enrichment of such pathways in the active subnetworks. For this type of analysis (over the 4 test cases) we use several curated databases of pathways. The association of the pathway to the disease can be either obvious from the pathway textual description, or it can be inferred by the presence of disease-related genes (at least two).

Figure 5(a) and (b) reports the number of affected pathways, obtained by combining the Reactome and DisGeNET information, that are enriched in the predicted active sub-networks. We note that only two methods (Core&Peel and ClustEx) are able to detect significant pathways on all the four test cases. ModuleDiscoverer finds at least one pathway in three test cases, while DEGs and KPM find pathways in two test cases and Degas in one only. Although Core&Peel and ClustEx do report enriched pathways on all four data sets, Core&Peel does find more such pathways (and for prostate cancer, substantially more).

For the two cancer datasets the number of detected pathways with at least two disease-associated genes is substantially larger than the number of the pathways detected for asthma and rheumatoid arthritis. This finding is consistent with the larger number of genes annotated on DisGeNET in cancer than asthma and rheumatoid arthritis, since the cancer is known to involve dysregulation on many cellular functions at once.

Although any of the competing methods (ClustEx, KPM, and ModuleDiscoverer, Degas) may perform better than Core&Peel in some particular case, Core&Peel is able to perform uniformly well across all four test cases, and moreover, it is the only method in the comparative evaluation that could find significant enrichment for many validated cancer pathways.

### 4.4 Relevant enriched pathways for four test cases

**Prostate cancer** : The majority of the pathways detected in prostate cancer are related to the regulation of the cell cycle and DNA repair, which are the main deregulated processes in cancer. In particular, one of the pathways that plays a role in the cell cycle is associated to the kinetochore, a protein that links the chromosome to microtubule and it is essential for proper chromosome segregation during mitosis. A study [51] showed that the expression of one kinetochore–associated protein was remarkably upregulated in prostate cancer. Knockdown of this protein repressed the ability of cell proliferation, migration, and invasion of prostate cancer cells. Another pathway with a role in the mitosis control is called *AURKA Activation by TPX2*. The protein TPX2 activetes Aurora A kinase (AURKA) which contributes to the regulation of cell cycle progression. It has been demonstrated [52] that overexpression of TPX2 improved proliferative, invasive and migratory abilities, and repressed apoptosis of the prostate cancer cells. Pathways involved in the DNA repair processes are for example *Resolution of D-loop Structures through Synthesis-Dependent Strand Annealing (SDSA)* and *HDR through Homologous Recombination (HRR)*. The SDSA and the Homology directed repair (HDR) are mechanisms in cells to repair double-strand DNA lesions. The SDSA is promoted by the DNA helicase and inactivating mutations in DNA helicase genes are frequently associated with various cancers. However, the overexpression of many DNA helicases is required for cancer cell proliferation or resistance to DNA damage [53]. In addition, mutations in genes that promote HDR are frequently observed in several cancers, including the prostate cancer [54]. *Cell-cell junction organization* and *Elastic fibre formation* pathways are also enriched. Elastic fiber are bundles of proteins found in the extracellular matrix. The extracellular matrix is commonly deregulated in cancer and affects the cancer progression, promoting metastasis [55]. Finally, *Fatty acids* pathway has been found to be associated to prostate and in fact several studies demonstrated the association of some fatty acids with prostate cancer risk [56, 57]. Moreover all the pathways enriched only in Core&Peel module include at least one gene with a disease-association in the DisGeNET database and/or DEGs. This highlights how these pathways can have a relevance in the prostate cancer.
**Colorectal cancer** : Similarly, all the pathways in colorectal cancer module include genes with disease-association and/or DEGs. Like prostate cancer case, pathways associated to the cell cycle have been found (*G2/M DNA replication checkpoint* and *Telomere C-strand synthesis initiation*). They are mainly involved during the DNA replication which represents a crucial point for the genome integrity. A deregulation of this process can lead an accumulation of genetic aberrations that promote diseases such as cancer [58]. Telomeres maintain genome integrity by protecting the end of the chromosome from deterioration. Telomere crisis can cause a wide array of genomic aberrations that can promote cancer progression [59]. Another characteristic of cancer is altered signal transduction which can lead to uninhibited growth. The *purinergic receptors* pathway is enriched in colorectal module, in fact it has been demonstrated that these receptors, after the binding with the ATP, affect tumor cell growth. In particular, it has been noticed an overexpression of some purienergic receptors in colon cancer [60], along with many malignant tumors. Another enriched pathway involved in the signal transduction is *G alpha signalling events*. Recent findings suggest that the prostaglandin E2, a proinflammatory product, stimulates colon cancer cell growth through a G protein–dependent signaling pathway [61]. Similary to prostate cancer, fatty acids metabolism is also involved in colorectal cancer. More precisely a rapid metabolism of *arachidonic acid* was reported in various stages of the malignancy, suggesting a possible link between dietary lipids and the incidence of colorectal cancer [62].
**Rheumatoid arthritis (RA)** : Rheumatoid arthritis is a chronic autoimmune disease that primarily affects the lining of the synovial joints. Among all the enriched pathways, two of them (*Neutrophil degranulation* and *Signaling by Interleukins*) are involved in the immune system and include both genes with disease-association (Figure 5b). Neutrophils are the most abundant leukocytes (white blood cells) in mammals and are one of the first-responders of inflammatory cells to migrate towards the site of inflammation. They contribute to RA pathology through the release of cytotoxic and immunoregulatory molecules, promoting the autoimmune processes that underly this disease [63]. The interleukins are proteins expressed by the immune system cells during an immune response. Different studies have demonstrated the involvement of interleukins in RA [64, 65]. In addition, several splice variants of interleukins have been discovered [66], highlighting the involvement of *mRNA splicing* pathway in RA. Other pathways detected only by Core&Peel have a role in immunity and inflammation such as *prefoldin* and *SUMO* proteins. Prefoldin is a family of proteins used in protein folding complexes and seems to function as proinflammatory signals [67]. SUMO is a small ubiquitin-like modifier involved in protein sumoylation, a post-translational-modification event. Sumoylation has been suggested to regulate multiple cellular processes, including inflammation [68].
**Asthma** : Asthma is a common long-term inflammatory disease of the airways of the lungs. In fact most of the enriched pathways identified only by Core&Peel are involved in the immune system. For example, the interaction between the *Toll-like receptors* (TLRs) and environmental allergens leads to release of various pro-inflammatory mediators from innate cells supporting asthma development [69]. Like in RA, different interleukins have a role in asthma pathogenesis [70, 71]. In addition, *butyrophilins* [72], *leukotrienes and eoxins* are molecules formed in response to inflammatory stimuli. In particular leukotrienes have a role in the pathophysiology of asthma including increased airway smooth muscle activity, microvascular permeability, and airway mucus secretion [73]. Another pathway enriched in asthma module and involved in the immune system is called *TNFs bind their physiological receptors*. TNFs function as cytokine and the binding with their specific receptors has crucial roles in both innate and adaptive immunity. In particular, TNF-*α* is a proinflammatory cytokine that has been implicated in many aspects of the airway pathology in asthma. Besides, preliminary studies have demonstrated an improvement in asthma quality of life and lung function in patients treated with anti–TNF-*α* therapy [74]. To conclude, a pathway identified only by Core&Peel which has four genes with disease-association is *G alpha signalling events*. G protein-coupled receptors (GPCRs) regulate numerous airway cell functions, and signaling events transduced by GPCRs are important in both asthma pathogenesis and therapy [75].

More details and figures on the enriched pathways for the four test cases are described in “ClueGO analysis” section in the Supplementary file.

### 4.5 Analysis of cellular transcriptional response to SARS-CoV-2 infection

We did re-analysis of three RNA-Seq case/control data sets on the human response to COVID-19 infection versus corresponding normal tissues/cell lines with several active sub-network detection algorithms. The analysis demonstrates a number of interesting functional modules that are enriched in these active networks (detected by one or more methods).

Functional annotations relative to Ebola infection is found enriched in tables S-8 (by all seven methods), S-6 and S-7, with strong p-values. This finding is of interest as it supports current attempts of re-positioning drugs developed to cure Ebola infections in patients affected by covid-19 (e.g. remdesivir: https://it.wikipedia.org/wiki/Remdesivir).

A second set of enriched functional modules is HIV-related (annotations in Tables S-8, S-6 and S-7). This observation also is consistent with the efforts in re-positioning of AIDS therapeutic agents for COVID-19 [76].

The *Staphylococcus aureus infection*, and *T-Cell antigen Receptor (TCR) pathway during Staphylococcus aureus infection WP3863* pathways are detected in Tables S-6, S-7 and S-8; while *Staphylococcus aureus infection* is detected in Table S-6. Staphylococcus aureus is known to produce pulmonary infections [77] thus it is a reasonable hypothesis that human response to S.aureus could share similarities at a molecular level with the COVID-19 response.

Cases of patients co-infected by COVID-19 and EBV (Epstein-Barr virus) have been reported [78], however our active subnetwork analysis suggests that also these two viruses may provoke similar response profiles in patients at a molecular level (EBV-related pathways are enriched in Tables S-6, S-7 and S-8).

Other pathogen-response modules enriched in the active subnetworks detected in all three data sets (Tables S-6, S-7 and S-8) are relative to Escherichia coli and human papillomavirus.

Two tables in suppl materials show enriched pathways for response to herpesvirus, Helicobacter pylori, Salmonella, cytomegalovirus, Yersinia, and Cholera.

The use of Anti-Herpesvirus Alkaloids in to cope with SARS-CoV-2 infection is advocated [79, 80], based on the similarity of respiratory symptoms induced by the bovine herpesvirus 1 in cattle, taking advantage of its inhibitory efficacy against SARS-CoV-2 replication. Cloroquine (Hydroxychloroquine) is a drug used in treatment of cholera [81] that is currently being assessed in the therapy of patients infected by COVID-19 [82, 83, 80].

We cannot rule out completely that some statistical associations found in the data could be explained by a case of co-infection by a variety of pathogens [84]. This event is however less likely for the data obtained through cellular lines, rather than patient samples. A second possibility is that the systemic human responses to these pathogens are greatly overlapping among themselves, besides being enriched in COVID-19 data, thus the statistical associations may not be independent of each other.

However, in a perspective of multi target drug design [85], it is important to be able to have wider choice of targets and drugs to consider based on the active sub-network analysis pre-screening.

In [32], using DEA analysis among other analytic tools, it is noted that infection by COVID-19 has a molecular fingerprint charachterizd by “exuberant inflammatory cytokine production as a defining and driving feature of COVID-19”, ad this is noted also in [31]. Our analysis of active subnetworks confirms this finding at the level of functional modules in pathways: *positive regulation of cytokine production involved in inflammatory response* (by Core&Peel), *regulation of cytokine production involved in inflammatory response* (by Core&Peel), *cytokine production involved in inflammatory response* (by Core&Peel), and *Cytokines and Inflammatory Response WP530* (by Core&Peel and ClustEx) in Table S-8. Pathway *Cytokines and Inflammatory Response WP530* is found enriched also in Table S-6 and S-7.

Also in [32] it is in noted a high expression of IL-6 as part of the characteristic COVID-19 signature. The importance of the Interleukin activation is also shown in our analysis, as the pathway *IL-10 Anti-inflammatory Signaling Pathway WP4495*, which has regulatory effects on IL-6, is found enriched in Tables S-6 and S-7, by ClustEx, and in Table S-8, by Core&Peel.

Transient receptor potential channels have been studied for their role in inflammatory processes and as therapeutic targets ([86]. [87]. [88]. [89]). We could find the pathway *Inflammatory mediator regulation of TRP channels* enriched in Tables S-8 and S-6. This observation supports another possible line in the search for COVID-19 treatments [90, 91].

Consistently with known mechanisms involved in SARS-CoV-2 infection progression [92], we have found several apoptosis-related and lymphocite-related pathways in all three data sets.

Current research on network-based analysis of COVID-19 focuses mainly in analyzing hostpathogen protein interaction maps [93, 94, 95] in order to find druggable target interactions. A second approach by Gysi et al. [96] uses host-pathogens protein interaction to extract a sub-network of a large human PPI network, which is further analysed for possible drug targets. Here we change focus, analyzing the modularity and enrichment of active sub-network in the host’s response to the infection within a global host co-expression network.

### 4.6 Limitations

In our active sub-network detection approach we have as input both a biological gene interaction network (called the “bio-network” data) and a list of gene expression data on cohorts of cases and controls for a disease under study (called the “perturbation” data). The bio-network represents a global and essentially static picture of the cell’s machinery, while the perturbation represents the dynamic element we wish to study within the context of the bio-network. All the methods we compare with imply these two roles (static/dynamic) for the two input (network/perturbation), although they may differ in many other aspects.

A limitation of this view on the data, is that the phenomenon of network rewiring in response to the stimulus is captured rather imperfectly. Rare novel gene interactions (i.e. new edges in the network) arising as the result of the stimulus are not included in this model, and cannot be predicted.

There is a different family of approaches (see e.g. [97]) that focus instead on the network rewiring phenomenon, by a more direct experimental assessment of dynamics of physical and genetic networks, through experimental mapping of networks. In this setting typically one would produce and analyze two full networks, one representing the status of the system before and one after the stimulus, in order to capture the variations. Here there is a different trade-off between the complexity of the experimental set up and the depth of the downstream ‘in silico’ analysis one can employ.

### 4.7 Conclusions

In conclusion, we have shown the state-of-the-art performance of the Core&Peel method for the task of finding functional modules and active functional sub-networks in large co-expression networks. Comparison with several active subnetwork detection algorithms on four benchmark test cases show that Core&Peel outperforms each competing method on some data set, and has more uniform performance across all the four benchmark data sets. As we do not believe in this area there is a ‘silver bullet’, we apply Core&Peel along with several other methods on COVID19-related transcriptomic data, recently made available. The combined output of these algorithms is able to find many significantly enriched active pathways that are at the center of current research efforts, as well as pointing at possible new directions of research for drug re-positioning.

## 5 Acknowledgements

We wish to thank Prof. Monica Bianchini for her help and support in conducting this research.

## Footnotes

1 The only methods mentioned in both lists are SPICi and ClusterOne, which are used in the functional module detection DREAM challenge as part of ensemble or hybrid computational pipelines.

2 Similar results if we take into account hits and spread hubs without any Jaccard filtering (data not shown).

