## Supplementary-Materials for "Finding disease modules for cancer and COVID-19 in gene co-expression networks with the Core&Peel method": supplementary.pdf

M. Lucchetta* M. Pellegrini†

May 27, 2020

*IIT-CNR, Pisa, Italy, and Dipartimento di Biotecnologie, Chimica e Farmacia, Università degli Studi di Siena, Siena, Italy

†IIT-CNR, Pisa, Italy. Contact author:

### 1 Supplementary Figures

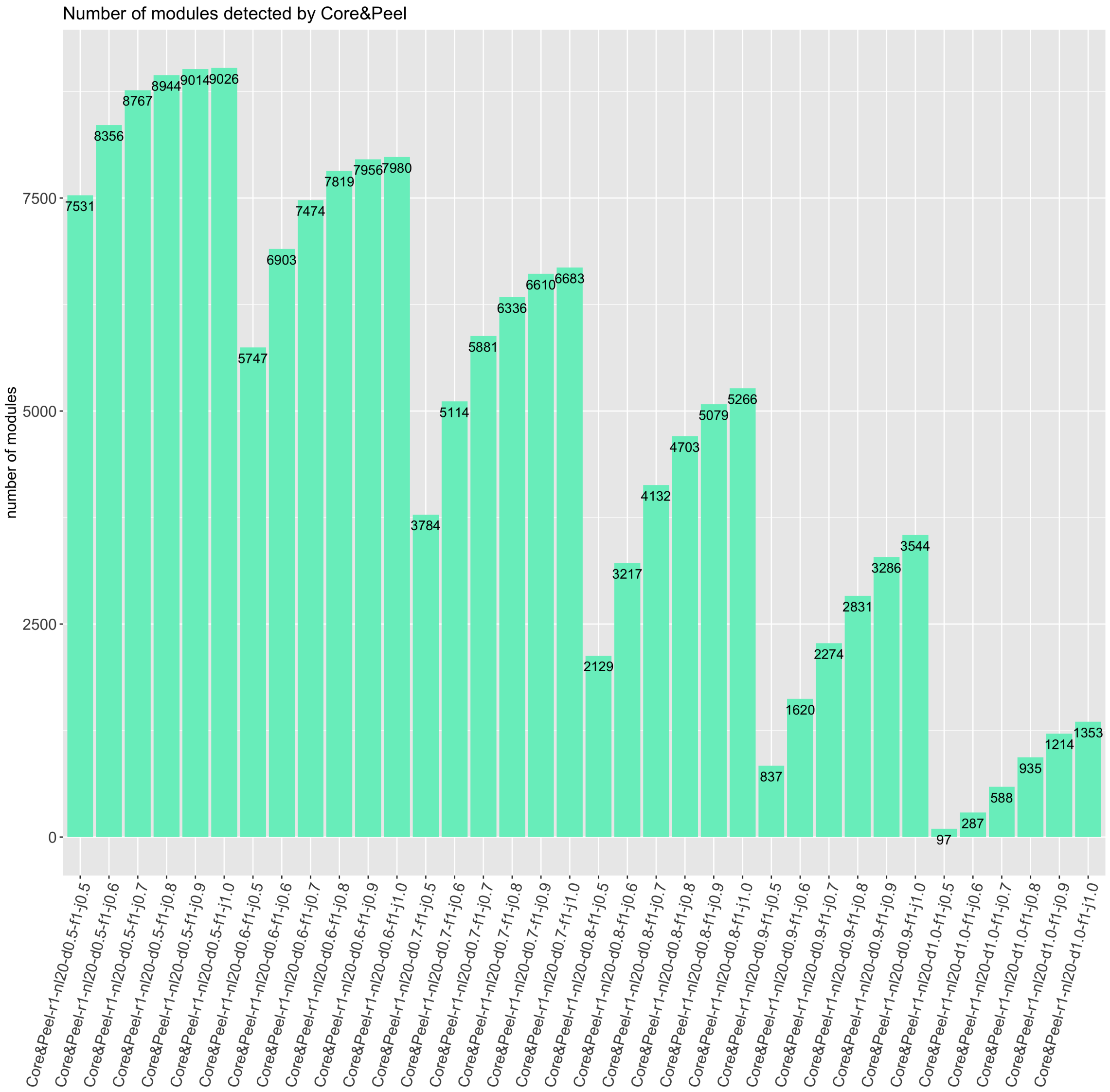

Figure S 1: Number of modules detected by Core&Peel according to different parameters.

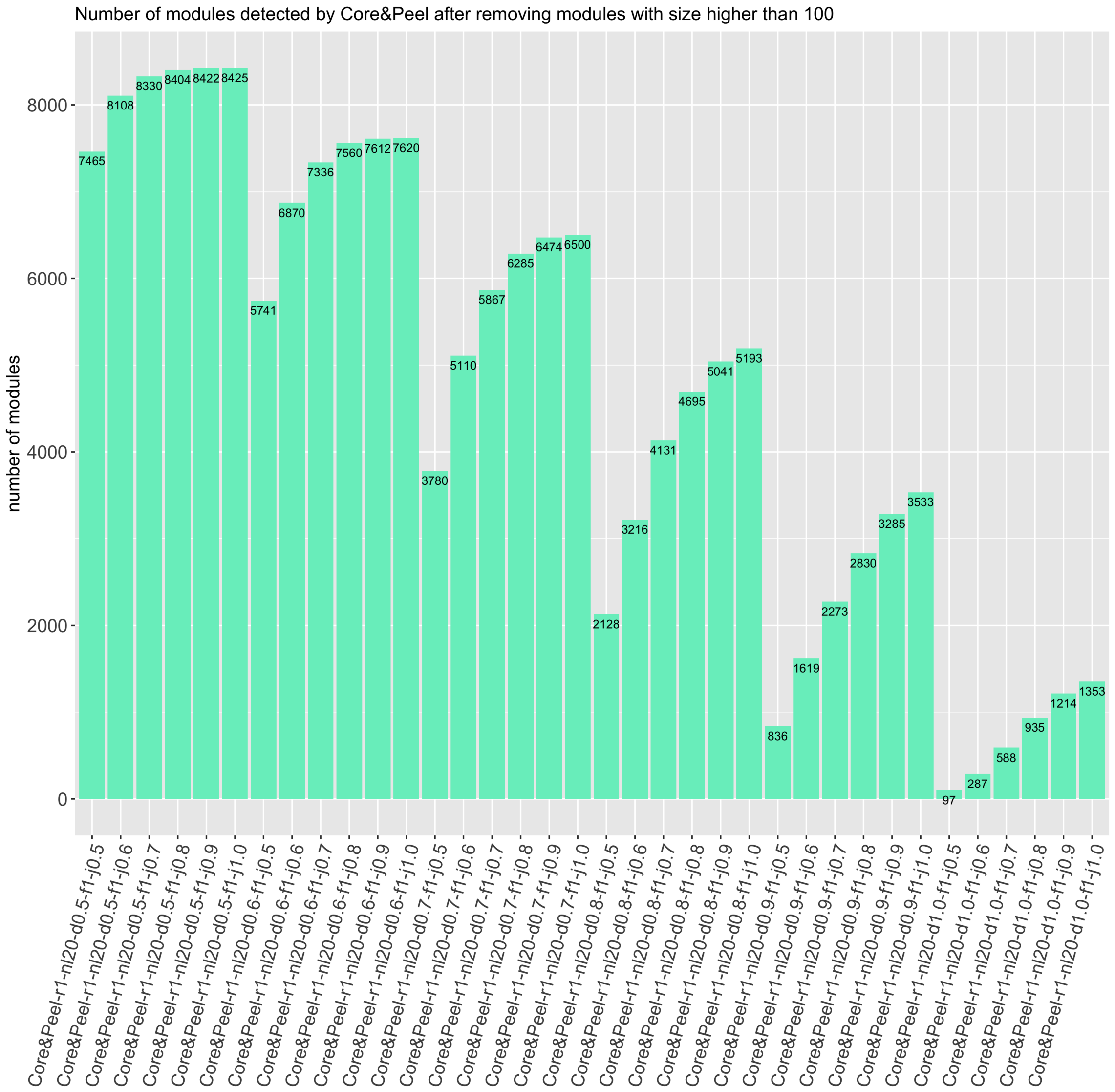

Figure S 2: Number of modules detected by Core&Peel after removing modules with size higher than 100.

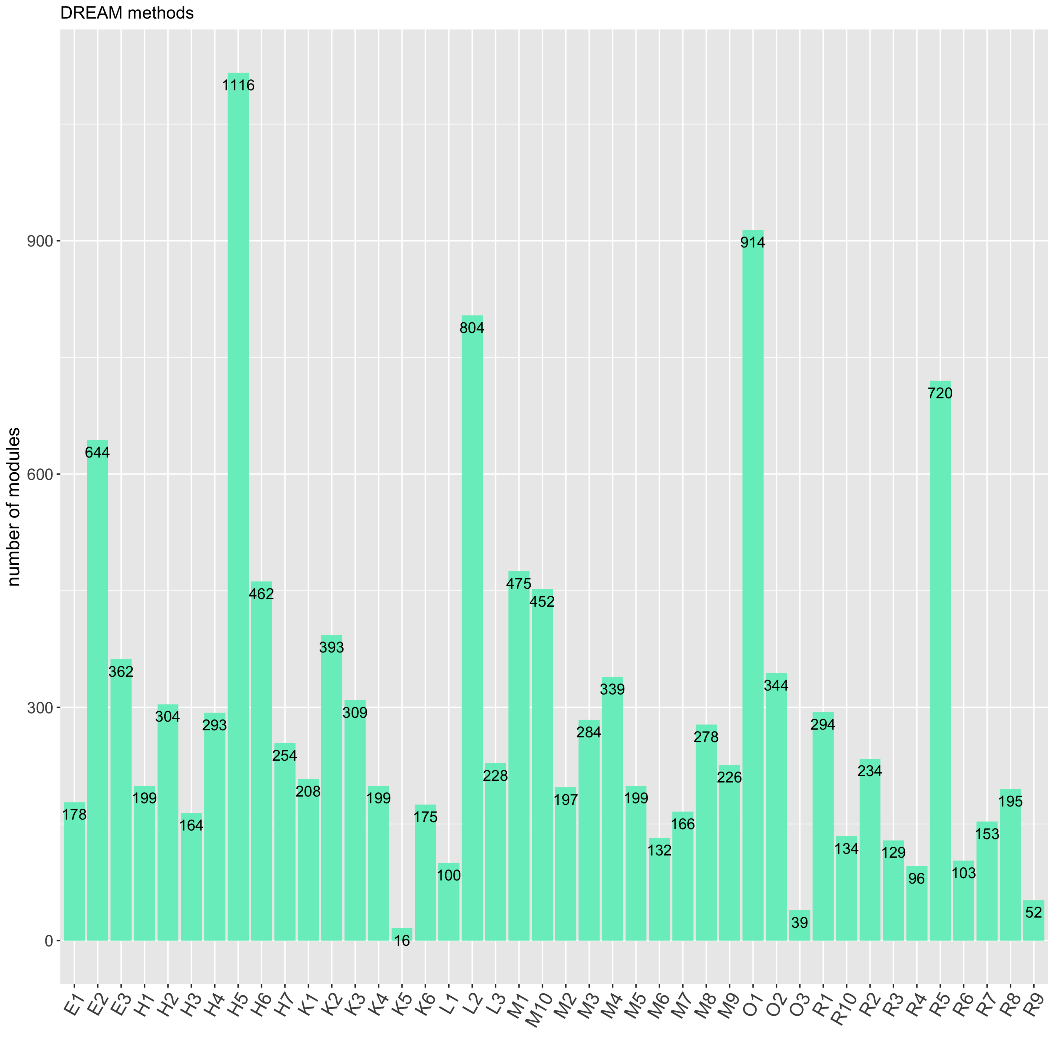

Figure S 3: Number of modules detected by the DREAM methods.

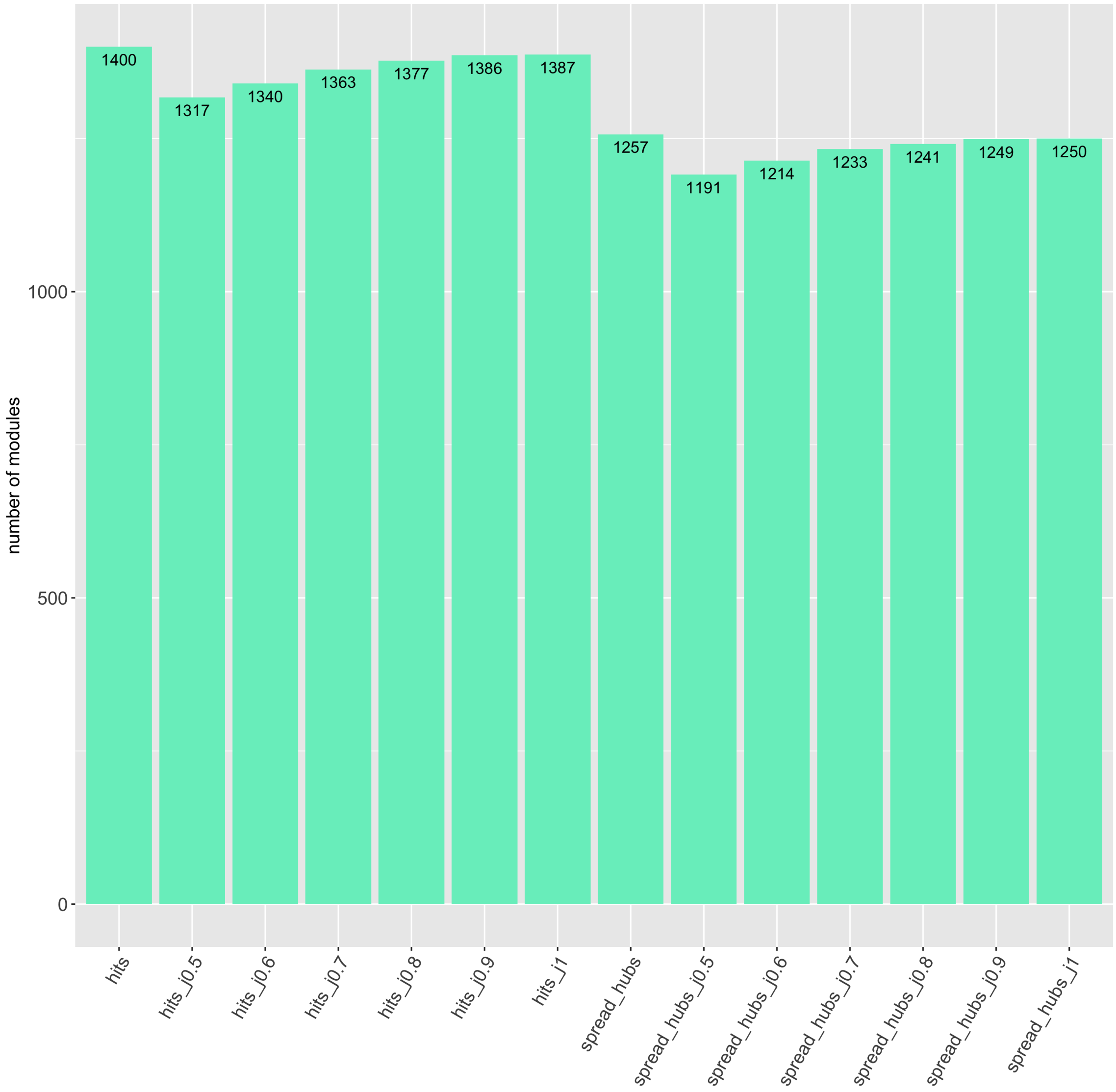

Figure S 4: Number of modules detected by further method which detects disease modules with overlap. Hits and spread_hubs have not been filtered for Jaccard coefficient.

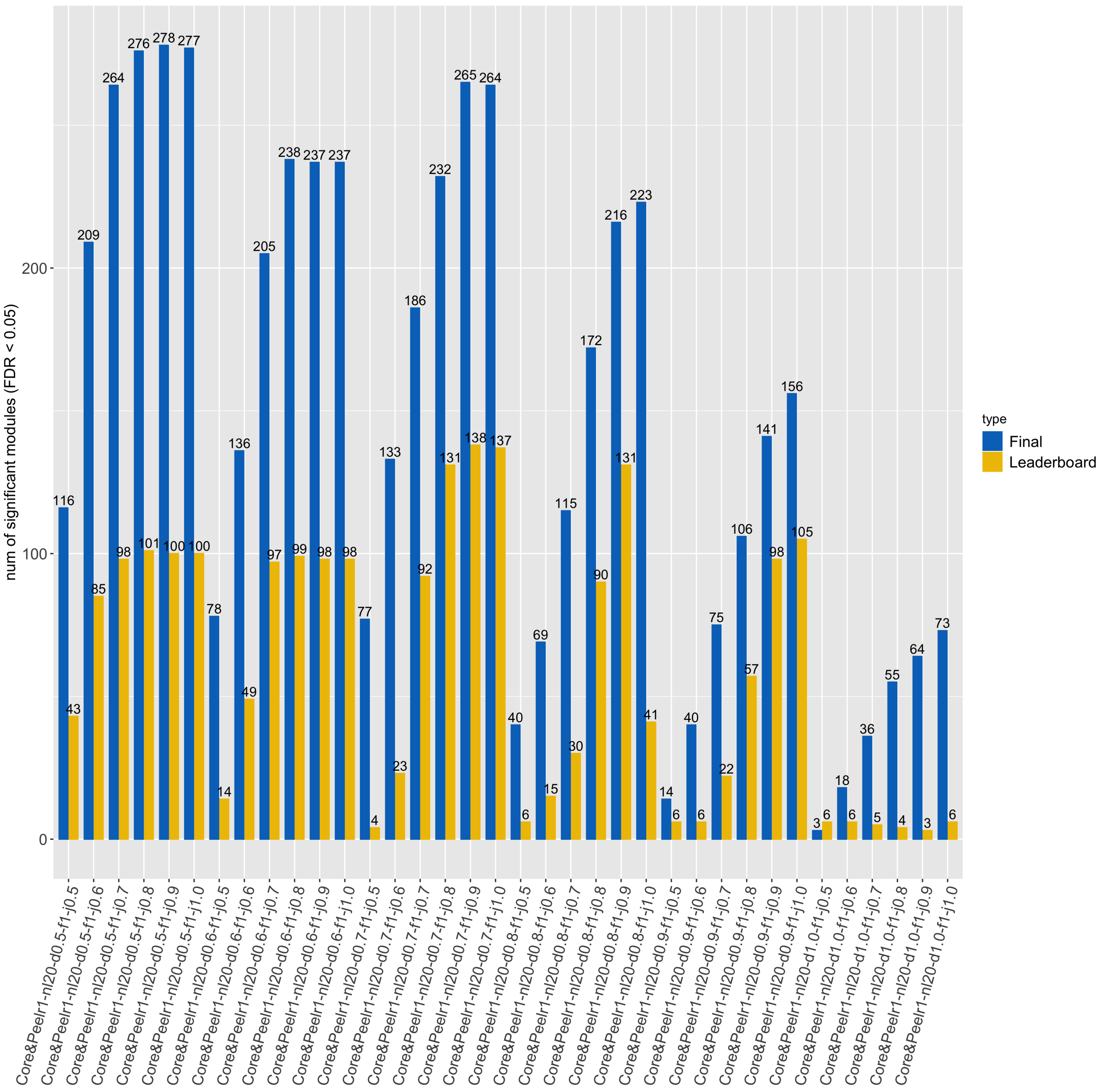

Figure S 5: Number of enriched disease modules detected by Core&Peel before removing modules with size higher than 100.

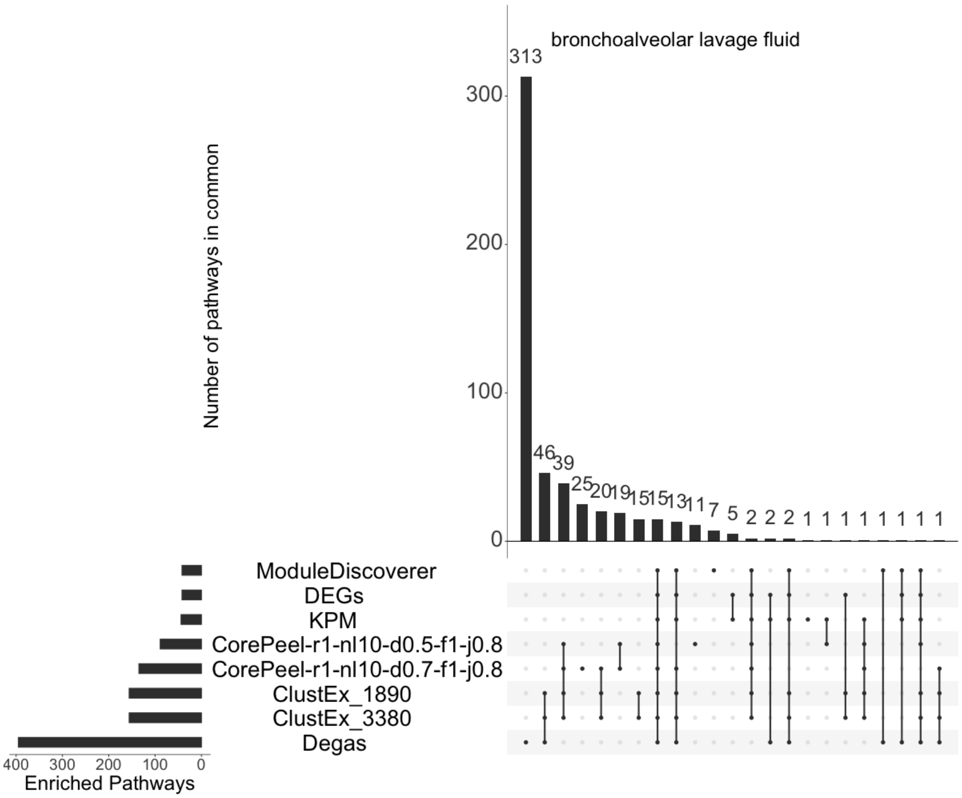

Figure S 6: Overlap of Reactome enriched pathways among all method combinations in BALF case study.

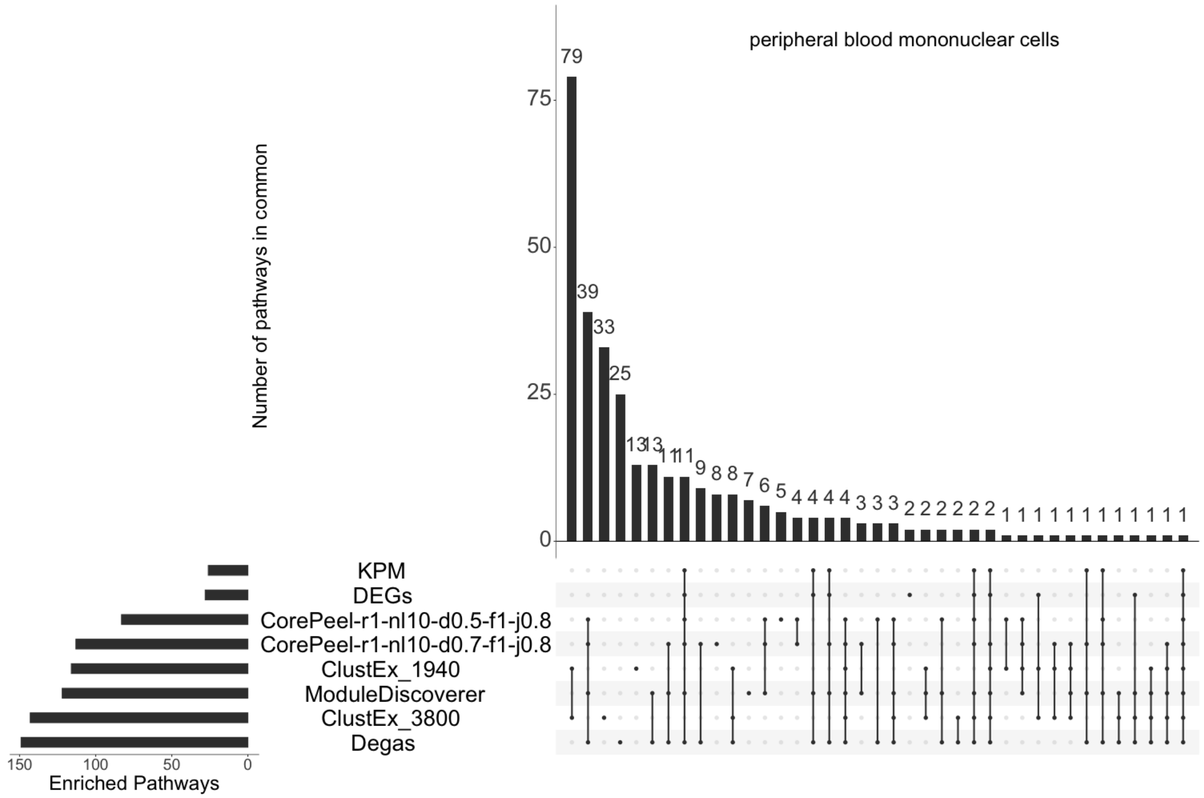

Figure S 7: Overlap of Reactome enriched pathways among all method combinations in PBMC case study.

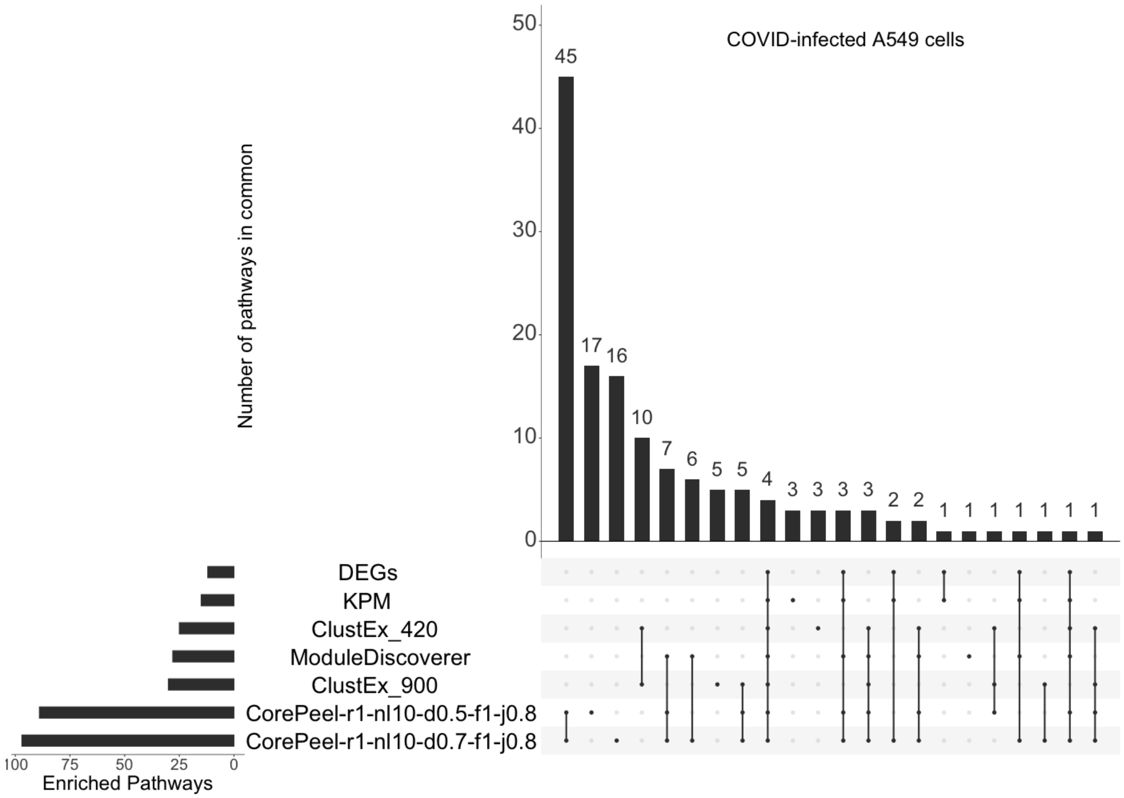

Figure S 8: Overlap of Reactome enriched pathways among all method combinations in COVID-infected cells case study.

### 2 Supplementary Tables

| | exp(-2) | exp(-3) | exp(-4) | exp(-5) | exp(-6) | exp(-7) |
|---|---|---|---|---|---|---|
| Core&Peel-r1-nl20-d0.7-f1-j0.8-M6 | 132 | 103 | 64 | 56 | 47 | 43 |
| M6 | 132 | 132 | 107 | 72 | 61 | 54 |
| Core&Peel-r1-nl20-d0.7-f1-j0.8-L2 | **804** | 673 | **449** | **354** | **278** | **230** |
| L2 | 800 | 708 | 299 | 155 | 92 | 63 |
| Core&Peel-r1-nl20-d0.7-f1-j0.8-K2 | 393 | 318 | **201** | **168** | **137** | **117** |
| K2 | 393 | 347 | 165 | 100 | 71 | 51 |
| Core&Peel-r1-nl20-d0.7-f1-j0.8-K5 | 16 | 14 | 9 | 7 | **7** | **7** |
| K5 | 16 | 16 | 12 | 7 | 3 | 2 |
| Core&Peel-r1-nl20-d0.7-f1-j0.8-M2 | 197 | 154 | 97 | 84 | 68 | **60** |
| M2 | 197 | 186 | 139 | 89 | 68 | 57 |
| Core&Peel-r1-nl20-d0.7-f1-j0.8-E1 | 178 | 137 | 84 | 73 | 58 | **54** |
| E1 | 178 | 172 | 130 | 94 | 64 | 52 |
| Core&Peel-r1-nl20-d0.7-f1-j0.8-M9 | 226 | 180 | **108** | **94** | **75** | **66** |
| M9 | 226 | 205 | 103 | 66 | 53 | 46 |
| Core&Peel-r1-nl20-d0.7-f1-j0.8-R6 | **103** | **80** | **51** | **45** | **38** | **37** |
| R6 | 102 | 87 | 38 | 31 | 23 | 21 |
| Core&Peel-r1-nl20-d0.7-f1-j0.8-M7 | 166 | 130 | 81 | 70 | **55** | **51** |
| M7 | 166 | 160 | 108 | 72 | 49 | 32 |
| Core&Peel-r1-nl20-d0.7-f1-j0.8-O2 | **344** | **282** | **176** | **148** | **119** | **102** |
| O2 | 337 | 281 | 34 | 17 | 10 | 7 |
| Core&Peel-r1-nl20-d0.7-f1-j0.8-H7 | 254 | **203** | **126** | **107** | **86** | **75** |
| H7 | 254 | 189 | 1 | 0 | 0 | 0 |
| Core&Peel-r1-nl20-d0.7-f1-j0.8-R3 | **129** | 100 | 61 | 53 | 43 | 41 |
| R3 | 126 | 114 | 74 | 59 | 52 | 42 |
| Core&Peel-r1-nl20-d0.7-f1-j0.8-K4 | 199 | 157 | 98 | **85** | **69** | **61** |
| K4 | 199 | 188 | 124 | 71 | 56 | 50 |
| Core&Peel-r1-nl20-d0.7-f1-j0.8-K3 | **309** | 251 | **159** | **136** | **108** | **93** |
| K3 | 308 | 278 | 150 | 87 | 64 | 49 |
| Core&Peel-r1-nl20-d0.7-f1-j0.8-M5 | **199** | 157 | 98 | **85** | **69** | **61** |
| M5 | 198 | 185 | 124 | 81 | 55 | 42 |
| Core&Peel-r1-nl20-d0.7-f1-j0.8-R7 | 153 | 119 | **74** | **63** | **51** | **47** |
| R7 | 153 | 138 | 64 | 44 | 33 | 28 |
| Core&Peel-r1-nl20-d0.7-f1-j0.8-R2 | 234 | 187 | 114 | **99** | **79** | **69** |
| R2 | 234 | 217 | 145 | 82 | 58 | 47 |
| Core&Peel-r1-nl20-d0.7-f1-j0.8-H6 | **462** | 377 | **241** | **200** | **161** | **139** |
| H6 | 461 | 391 | 56 | 18 | 9 | 7 |
| Core&Peel-r1-nl20-d0.7-f1-j0.8-R10 | 134 | **105** | **66** | **57** | **48** | **44** |
| R10 | 134 | 94 | 1 | 0 | 0 | 0 |
| Core&Peel-r1-nl20-d0.7-f1-j0.8-H5 | **1116** | 937 | **626** | **491** | **377** | **305** |
| H5 | 1110 | 952 | 201 | 114 | 79 | 59 |
| Core&Peel-r1-nl20-d0.7-f1-j0.8-H4 | 293 | 238 | 151 | **130** | **103** | **88** |
| H4 | 293 | 269 | 180 | 91 | 67 | 38 |
| Core&Peel-r1-nl20-d0.7-f1-j0.8-R4 | 96 | 75 | 47 | 41 | 35 | 34 |
| R4 | 96 | 88 | 58 | 44 | 37 | 34 |
| Core&Peel-r1-nl20-d0.7-f1-j0.8-M10 | **452** | 368 | **235** | **195** | **156** | **134** |
| M10 | 451 | 414 | 154 | 71 | 56 | 34 |
| Core&Peel-r1-nl20-d0.7-f1-j0.8-M4 | 339 | 279 | 176 | **148** | **119** | **102** |

| | | | | | | |
|---|---|---|---|---|---|---|
| M4 | 339 | 310 | 187 | 114 | 80 | 63 |
| Core&Peel-r1-nl20-d0.7-f1-j0.8-H1 | 199 | 157 | 98 | 85 | 69 | 61 |
| H1 | 199 | 189 | 149 | 107 | 74 | 57 |
| Core&Peel-r1-nl20-d0.7-f1-j0.8-R5 | 720 | 604 | **405** | **323** | **252** | **213** |
| R5 | 720 | 652 | 227 | 130 | 77 | 63 |
| Core&Peel-r1-nl20-d0.7-f1-j0.8-O1 | 914 | 766 | **514** | **407** | **316** | **257** |
| O1 | 914 | 799 | 116 | 62 | 41 | 34 |
| Core&Peel-r1-nl20-d0.7-f1-j0.8-O3 | 39 | **35** | **20** | **18** | **16** | **16** |
| O3 | 39 | 27 | 0 | 0 | 0 | 0 |
| Core&Peel-r1-nl20-d0.7-f1-j0.8-H3 | 164 | 128 | 79 | 68 | **54** | **50** |
| H3 | 164 | 159 | 115 | 77 | 51 | 39 |
| Core&Peel-r1-nl20-d0.7-f1-j0.8-M3 | 284 | 230 | 144 | **123** | **98** | **84** |
| M3 | 284 | 265 | 180 | 100 | 75 | 62 |
| Core&Peel-r1-nl20-d0.7-f1-j0.8-L1 | **100** | 77 | 48 | 42 | 36 | 35 |
| L1 | 99 | 90 | 62 | 54 | 49 | 40 |
| Core&Peel-r1-nl20-d0.7-f1-j0.8-K1 | 208 | 165 | 101 | 88 | 70 | **63** |
| K1 | 208 | 201 | 154 | 90 | 70 | 56 |
| Core&Peel-r1-nl20-d0.7-f1-j0.8-R1 | 294 | 238 | 151 | **130** | **103** | **88** |
| R1 | 294 | 264 | 175 | 111 | 84 | 67 |
| Core&Peel-r1-nl20-d0.7-f1-j0.8-M1 | 475 | 390 | **248** | **207** | **167** | **145** |
| M1 | 474 | 416 | 167 | 97 | 78 | 61 |
| Core&Peel-r1-nl20-d0.7-f1-j0.8-H2 | **304** | 246 | 156 | **134** | **106** | **91** |
| H2 | 303 | 280 | 182 | 127 | 92 | 67 |
| Core&Peel-r1-nl20-d0.7-f1-j0.8-E3 | 362 | 296 | **188** | **157** | **127** | **108** |
| E3 | 362 | 315 | 109 | 52 | 35 | 22 |
| Core&Peel-r1-nl20-d0.7-f1-j0.8-E2 | **644** | 537 | **352** | **286** | **226** | **194** |
| E2 | 643 | 580 | 232 | 126 | 77 | 60 |
| Core&Peel-r1-nl20-d0.7-f1-j0.8-L3 | 228 | 182 | 111 | 96 | **77** | **68** |
| L3 | 228 | 216 | 126 | 94 | 71 | 57 |
| Core&Peel-r1-nl20-d0.7-f1-j0.8-K6 | **175** | **134** | **82** | **71** | **56** | **52** |
| K6 | 173 | 130 | 39 | 21 | 12 | 9 |
| Core&Peel-r1-nl20-d0.7-f1-j0.8-R8 | **195** | 152 | **96** | **83** | **66** | **60** |
| R8 | 194 | 166 | 83 | 56 | 45 | 38 |
| Core&Peel-r1-nl20-d0.7-f1-j0.8-R9 | 52 | 44 | 25 | **23** | **20** | **19** |
| R9 | 52 | 46 | 25 | 18 | 13 | 10 |
| Core&Peel-r1-nl20-d0.7-f1-j0.8-M8 | 278 | 223 | 140 | **119** | **96** | **82** |
| M8 | 278 | 245 | 164 | 104 | 70 | 54 |
| mean Core&Peel | **299.1** | 243.5 | **156** | **129.7** | **103.4** | **89.2** |
| mean DREAM | 298.4 | 265.7 | 118.1 | 71.5 | 51.3 | 40 |

Table S 1: Summary of the GO enrichment analysis results. The values shown represent the number of predicted modules with GO enrichment fdr below different thresholds. Core&Peel-r1-nl20-d0.7-f1-j0.8 was compared with DREAM methods after applying the CRank algorithm to Core&Peel in order to get the same number of modules. When Core&Peel detects more enriched modules than DREAM methods the corresponding number is in bold. The last row represents the mean of the enriched modules for Core&Peel and DREAM methods for each threshold.

| | exp(-2) | exp(-3) | exp(-4) | exp(-5) | exp(-6) | exp(-7) |
|---|---|---|---|---|---|---|
| Core&Peel-r1-nl20-d0.7-f1-j0.8 | **6285** | **5332** | **3343** | **2169** | **1445** | **1063** |
| hits jaccard0.8 | 1377 | 953 | 346 | 189 | 111 | 88 |
| hits | 1400 | 975 | 362 | 206 | 117 | 95 |
| spread hubs jaccard0.8 | 1241 | 854 | 329 | 169 | 108 | 87 |
| spread hubs | 1257 | 870 | 345 | 183 | 114 | 91 |

Table S 2: Summary of the GO enrichment analysis results comparing Core&Peel-r1-nl20-d0.7-f1-j0.8 with the method which detects overlapping communities. Hits jaccard0.8 and spread hubs jaccard0.8 are filtered in order to have the maximum Jaccard separation equal to 0.8, hits and spread hubs are the same methods but without any filtering. The numbers in bold highlight the best method in detecting more significant GO terms.

| | exp(-2) | exp(-3) | exp(-4) | exp(-5) | exp(-6) | exp(-7) |
|---|---|---|---|---|---|---|
| Core&Peel-r1-nl20-d0.7-f1-j0.8 | 1 | **0.848** | **0.532** | **0.345** | **0.230** | **0.169** |
| hits jaccard0.8 | 1 | 0.692 | 0.251 | 0.137 | 0.08 | 0.064 |
| hits | 1 | 0.696 | 0.258 | 0.147 | 0.083 | 0.068 |
| spread hubs jaccard0.8 | 1 | 0.688 | 0.265 | 0.136 | 0.087 | 0.07 |
| spread hubs | 1 | 0.692 | 0.274 | 0.145 | 0.09 | 0.072 |

Table S 3: Fraction of modules with GO enrichment fdr below different thresholds across all the modules detected as significant (fdr $\leq$ 0.05). Again, the numbers in bold highlight the best method in detecting significant GO terms.

| | case | control | DEGs |
|---|---|---|---|
| Prostate cancer | 478 | 52 | 1301 |
| Asthma | 12 | 15 | 100 |
| Rheumatoid | 18 | 15 | 21 |
| Colorectal cancer | 451 | 41 | 2995 |
| BALF | 2 | 3 | 630 |
| PBMC | 3 | 3 | 538 |
| COVID-cells | not specified | not specified | 38 |

Table S 4: case: number of samples with disease, control: number of healthy samples, DEGs: number of differentially expressed genes included in the network.

| | BALF | PBMC | COVID-cells |
|---|---|---|---|
| Core&Peel-r1-nl10-d0.5-f1-j0.8 | 3380 | 3809 | 904 |
| Core&Peel-r1-nl10-d0.7-f1-j0.8 | 1895 | 1943 | 423 |
| MD | 1183 | 1329 | 273 |
| Degas | 4112 | 122 | - |
| ClustEx | 3380-1890 | 3800-1940 | 900-420 |
| KPM | 605 | 520 | 38 |

Table S 5: Number of genes in each active sub-network in the three COVID-19 cases. Both Core&Peel configurations have been tested. Two versions of ClustEx have been generated in order to get modules with size comparable with Core&Peel. It was not possible to run Degas in COVID-cells case since the expression matrix was not available.

| **Pathway** | **Core&Peel d0.5-j0.8** | **Core&Peel d0.7-j0.8** | **MD** | **ClustEx 3380** | **ClustEx 1890** | **KPM** | **Degas** | **DEGs** |
|---|---|---|---|---|---|---|---|---|
| Staphylococcus aureus infection | 1.42 | NA | NA | NA | NA | NA | NA | NA |
| Inflammatory mediator regulation of TRP channels | 1.31 | NA | NA | NA | NA | NA | NA | NA |
| Cytokines and Inflammatory Response WP530 | NA | NA | 1.35 | 1.68 | 2.0 | NA | NA | NA |
| Ebola Virus Pathway on Host WP4217 | NA | NA | 1.90 | 6.02 | 5.0 | NA | 6.48 | NA |
| Human papillomavirus infection | NA | NA | NA | 2.38 | 4.2 | NA | NA | NA |
| Vibrio cholerae infection | NA | NA | NA | 2.38 | 2.63 | NA | 1.88 | NA |
| Epstein-Barr virus infection | NA | NA | NA | 2.12 | 1.84 | NA | 3.88 | NA |
| Kaposi sarcoma-associated herpesvirus infection | NA | NA | NA | 1.34 | 3.28 | NA | 2.44 | NA |
| Human cytomegalovirus infection | NA | NA | NA | 1.33 | 3.10 | NA | 1.96 | NA |
| Epithelial cell signaling in Helicobacter pylori infection | NA | NA | NA | NA | 2.53 | NA | 2.87 | NA |
| Yersinia infection | NA | NA | NA | NA | 2.39 | NA | 2.67 | NA |
| Human immunodeficiency virus 1 infection | NA | NA | NA | NA | 1.95 | NA | 1.47 | NA |
| Salmonella infection | NA | NA | NA | NA | 1.81 | NA | 6.57 | NA |
| T-Cell antigen Receptor (TCR) pathwayduring Staphylococcus aureus infection WP3863 | NA | NA | NA | 3.08 | 2.89 | NA | 4.19 | NA |
| Pathogenic Escherichia coli infection | NA | NA | NA | NA | NA | NA | 1.94 | NA |
| IL-10 Anti-inflammatory Signaling Pathway WP4495 | NA | NA | NA | NA | NA | NA | 1.33 | NA |

Table S 6: Most interesting enriched pathways in BALF-COVID19 case. The negative adjusted p-values in log10-scale are reported. If one pathway is not significant, NA is reported. Both Core&Peel configurations (Core&Peel-r1-nl10-d0.5-f1-j0.8 and Core&Peel-r1-nl10-d0.7-f1-j0.8) have been tested. Consequently two versions of ClustEx have been generated in order to get them comparable with the two Core&Peel configurations according to the different module sizes. MD: ModuleDiscoverer; KPM: KeyPathwayMiner; DEGs: Differentially Expressed Genes.

| **Pathways** | **Core&Peel d0.5-j0.8** | **Core&Peel d0.7-j0.8** | **MD** | **KPM** | **ClustEx 3800** | **ClustEx 1940** | **Degas** | **DEGs** |
|---|---|---|---|---|---|---|---|---|
| Staphylococcus aureus infection | NA | 2.60 | NA | NA | NA | 1.74 | NA | NA |
| Human immunodeficiency virus infectious disease | 5.72 | 5.82 | 3.87 | 1.49 | NA | NA | NA | NA |
| Inflammatory Response Pathway WP453 | 3.89 | 5.16 | NA | NA | 1.66 | 1.45 | NA | NA |
| T-Cell antigen Receptor (TCR) pathway during Staphylococcus aureus infection WP3863 | 2.53 | NA | NA | NA | 2.75 | 1.48 | NA | NA |
| Pathogenic Escherichia coli infection WP2272 | 2.18 | NA | NA | NA | NA | NA | NA | NA |
| Ebola Virus Pathway on Host WP4217 | 1.49 | NA | NA | NA | 5.77 | 4.64 | NA | NA |
| Epstein-Barr virus infection | NA | NA | NA | NA | 3.19 | 2.53 | NA | NA |
| Vibrio cholerae infection | NA | NA | NA | NA | NA | 2.02 | NA | NA |
| Epithelial cell signaling in Helicobacter pylori infection | NA | NA | NA | NA | NA | 1.37 | NA | NA |
| Human papillomavirus infection | NA | NA | NA | NA | 2.29 | NA | NA | NA |
| Cytokines and Inflammatory Response WP530 | NA | NA | NA | NA | 3.80 | 1.77 | NA | NA |
| IL-10 Anti-inflammatory Signaling Pathway WP4495 | NA | NA | NA | NA | 1.38 | NA | NA | NA |

Table S 7: Most interesting enriched pathways in PBMC-COVID19 case. The negative adjusted p-values in log10-scale are reported. If one pathway is not significant, NA is reported. Both Core&Peel configurations (Core&Peel-r1-nl10-d0.5-f1-j0.8 and Core&Peel-r1-nl10-d0.7-f1-j0.8) have been tested. Consequently two versions of ClustEx have been generated in order to get them comparable with the two Core&Peel configurations according to the different module sizes. MD: ModuleDiscoverer; KPM: KeyPathwayMiner; DEGs: Differentially Expressed Genes.

| Pathway | Core&Peel d0.5-j0.8 | Core&Peel d0.7-j0.8 | MD | ClustEx 900 | ClustEx 420 | KPM | DEGs |
|---|---|---|---|---|---|---|---|
| positive regulation of cytokine production involved in inflammatory response | 1.57 | 2.96 | NA | NA | NA | NA | NA |
| regulation of cytokine production involved in inflammatory response | 2.14 | 2.82 | NA | NA | NA | NA | NA |
| cytokine production involved in inflammatory response | 1.90 | 2.62 | NA | NA | NA | NA | NA |
| Inflammatory mediator regulation of TRP channels | 2.14 | 1.94 | NA | NA | NA | NA | NA |
| IL-10 Anti-inflammatory Signaling Pathway WP4495 | 2.87 | 2.87 | NA | NA | NA | NA | NA |
| Cytokines and Inflammatory Response WP530 | 1.49 | NA | NA | 2.8 | 1.62 | NA | NA |
| Recurrent Staphylococcus aureus infections | 1.42 | 2.72 | NA | NA | NA | NA | NA |
| Epstein-Barr virus infection | 10.16 | 13.69 | 6.18 | 6.70 | 7.84 | 7.33 | 7.33 |
| Human immunodeficiency virus 1 infection | 7.19 | 7.43 | 2.75 | 1.78 | 3.88 | NA | NA |
| Herpes simplex virus 1 infection | 4.60 | 7.28 | 4.31 | 3.32 | 4.05 | 5.73 | 5.73 |
| Staphylococcus aureus infection | 4.54 | 1.94 | 4.99 | 2.75 | NA | NA | NA |
| Human cytomegalovirus infection | 4.47 | 3.0 | 2.95 | NA | 1.38 | NA | NA |
| Salmonella infection | 2.23 | 3.29 | NA | NA | NA | NA | NA |
| Yersinia infection | 2.21 | NA | 1.82 | NA | NA | NA | NA |
| Human papillomavirus infection | 1.90 | 2.29 | 2.82 | NA | 1.37 | 2.81 | 2.81 |
| Pathogenic Escherichia coli infection | 1.75 | 2.77 | NA | NA | 1.74 | NA | NA |
| T-Cell antigen Receptor (TCR) pathway during Staphylococcus aureus infection WP3863 | 4.24 | 3.90 | 2.35 | NA | NA | NA | NA |
| Ebola Virus Pathway on Host WP4217 | 8.36 | 7.07 | 4.99 | 4.85 | 3.05 | 1.91 | 1.59 |

Table S 8: Most interesting enriched pathways in COVID19-cells case. The negative adjusted p-values in log10-scale are reported. If one pathway is not significant, NA is reported. Both Core&Peel configurations (Core&Peel-r1-nl10-d0.5-f1-j0.8 and Core&Peel-r1-nl10-d0.7-f1-j0.8) have been tested. Consequently two versions of ClustEx have been generated in order to get them comparable with the two Core&Peel configurations according to the different module sizes. MD: ModuleDiscoverer; KPM: KeyPathwayMiner; DEGs: Differentially Expressed Genes.

### 3 Supplementary Notes

#### 3.1 Algorithms for active subnetwork identification

We tested four existing active subnetwork identification algorithms. We selected the methods that take gene expression matrix and/or list of differentially expressed genes (DEGs) as their inputs [1], a part from the network (generally it is a protein-protein interaction network). In this case, we used the co-expression network by the DREAM challenge as input. The algorithms we used in the comparison are summarized as follows.

**ModuleDiscoverer** [2]: Starting from a network, the algorithm first approximates the underlying community structure by iterative enumeration of cliques from random seed nodes (one or more) in the network. Secondly, all the cliques are tested for their enrichment with the DEGs obtained from the gene expression data. Finally, significantly enriched cliques are assembled in a unique regulatory module taking each gene once.
We used the single-seed approach, which identifies cliques using only one seed node and the p-value cutoff was 0.01 (both default parameters). The p-value for each clique is calculated using a permutation-based test. To calculate this p-value we used the parameter of 20000 gene-sets.

**KeyPathwayMiner (KPM)** [3]: it detects highly connected subnetworks in which most genes show similar patterns of expression. Specifically, given network and gene expression data, KeyPathwayMiner identifies those maximal subgraphs where all but $k$ nodes of the subnetwork are expressed similarly in all but $l$ cases in the gene expression data. We provided an indicator flag to mark differentially expressed genes (0/1). We set k= 2 and l = 0 and INES strategy with GREEDY algorithm was performed. More modules were detected but most of them differs from one or two genes, so the best-scoring module was selected for further analysis.

**DEGAS** [4]: it identifies connected gene subnetworks significantly enriched for genes that are dysregulated in samples with disease. It takes the network and the gene expression matrix as inputs and each sample is labeled as case or control. We used default parameters a part from $l$ (the number of outlier cases) which was set as 20% of the case number (as suggested by the authors). The algorithm identified one regulatory module for case study, which were used for further analysis. Only in BALF and PBMC cases, it detected more than one module so we merged them together. Beside it was not possible to run Degas in COVID-cells case since the expression matrix was not available.

**ClustEx** [5]: the algorithm takes DEGs and network as input. The DEGs are clustered and partitioned into different groups by average linkage hierarchical clustering according to their distances in the network. In the extending step, the final modules are formed by adding intermediate genes on the k-shortest paths between the DEGs. The size of the largest module has to be chosen. The output consists of some small modules (about 2 or 3 genes for each) and one big module. We took the biggest module for the downstream analyses. The size of the largest module was chosen in order to be comparable with the module size of Core&Peel.

#### 3.2 Core&Peel: detection of active sub-network when nl = 20

Here we show the Core&Peel performance in detecting active subnetwork when nl = 20. We chose the Core&Peel-r1-nl20-d0.8-f1-j0.8 for the prostate and colorectal cancer cases, instead Core&Peel-r1-nl20-d0.5-f1-j0.8 in asthma and rheumatoid arthritis case. Core&Peel detected 743, 120, 33 and 837 modules in the prostate cancer, asthma, rheumatoid arthritis and colorectal cancer datasets, respectively. Since the significant modules were merged in one regulatory module, we checked if these modules were still significantly enriched with the DEGs; all the four modules got a p-value $< 10^{-12}$.

##### 3.2.1 Comparison with other active sub-network detection methods

We compared the Core&Peel modules with those detected by the four existing active subnetwork identification methods, namely ModuleDiscoverer (MD), KeyPathwayMiner (KPM), ClustEx and Degas. We also tested MD using the hypergeometric exact test (MD_hyperg) instead of the permutation-based test.
The number of genes in each module is reported in Table 9. Also in this case, regulatory modules of asthma and rheumatoid arthritis include a smaller number of genes than the prostate and colorectal cancer modules. Besides when we apply the hypergeometric test to MD, smaller modules have been identified than classic MD.

| | Prostate | Asthma | Colorectal | Rheumatoid arthritis |
|---|---|---|---|---|
| Core&Peel | 1679 | 1170 | 2220 | 416 |
| MD | 2708 | 192 | 2606 | 403 |
| MD_hyperg | 1435 | 173 | 1647 | 160 |
| KPM | 1266 | 74 | 2965 | 20 |
| ClustEx | 1700 | 1200 | 2300 | 350 |
| Degas | 122 | 37 | 44 | 85 |

Table 9: Number of genes in each active subnetwork detected by all the methods for the four case studies

Next, we compared the enriched pathways (Reactome-based) and plotted the overlap among all the method combinations (Figure 9). In this comparison we took into account the MD_hyperg. Globally, MD_hyperg, ClustEx and Core&Peel detected higher number of pathways than Degas, KPM and DEGs. In both cancer cases, Core&Peel and MD_hyperg share almost half of their pathways (around 60) and a large number of pathways was identified only by ClustEx (Figures 9A, 9B). In rheumatoid arthritis case, more than half of the pathways were shared by Core&Peel, ClustEx and MD_hyperg (Figure 9D) and there is a small overlap for the other combinations. Besides, a smaller number of enriched pathways were detected in asthma than the other three case studies and we can also notice there is a few overlap. In fact almost the entire set of enriched pathways of ClustEx is not in common with any other methods. Similarly, more than half of the pathways identified by Core&Peel, were not detected by the others (Figure 9C).

##### 3.2.2 Detection of disease-associated pathways

We calculated the number of enriched pathways with at least two genes associated to the specific disease (annotated on DisGeNET database). Also in this case we took into account MD_hyperg. The results are showed in Figure 10. Generally Core&Peel is able to identify a substantial number of pathways. In particular, it is able to detect the major number of pathways with disease-associated genes in colorectal cancer and asthma. Generally applying the hypergeometric test reduces the number of pathways with disease-associated-genes, more evident in the rheumatoid arthritis case (Figure 10B).

Finally, we conducted further enrichment analyses using different databases in order to investigate the enrichment with specific-disease pathways. The results are showed in Figure 11. The results for carcinoma and asthma pathways are similar to those obtained for nl = 10 (compare Figure 11 A and B with Figure in the main text). The Core&Peel modules are the unique ones to be significantly enriched for colon and prostate cancer specific pathways such as neoplasm of the colon (Figure 11C) and abnormal prostate morphology, prostate cancer and prostate neoplasm (Figure 11D). This highlights how Core&Peel (with nl = 20) was able to find more cancer type-specific modules than the others algorithms. Again no significantly enriched pathways were found for rheumatoid arthritis

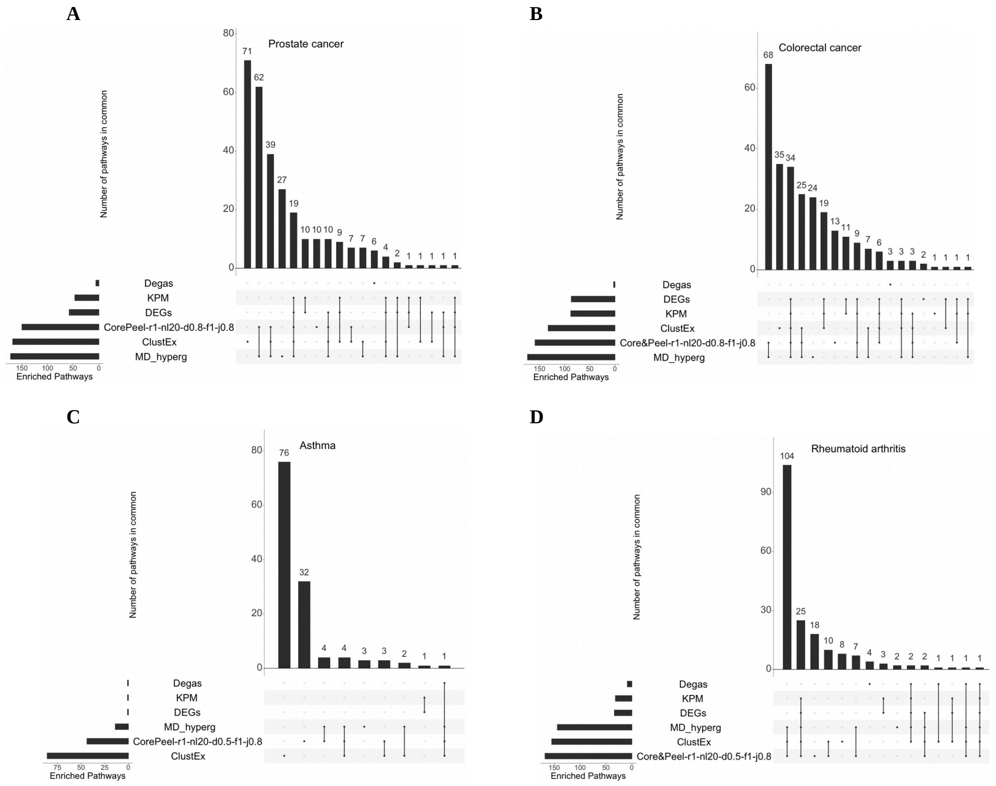

Figure 9: Overlap of enriched pathways among all method combinations in four study cases. The bars on the bottom-left represent the number of enriched pathways detected by the methods. The bars on the main plot represent the number of enriched pathways in common between methods marked with black points on the panel below. The plot was generated by the R package *UpSetR*.

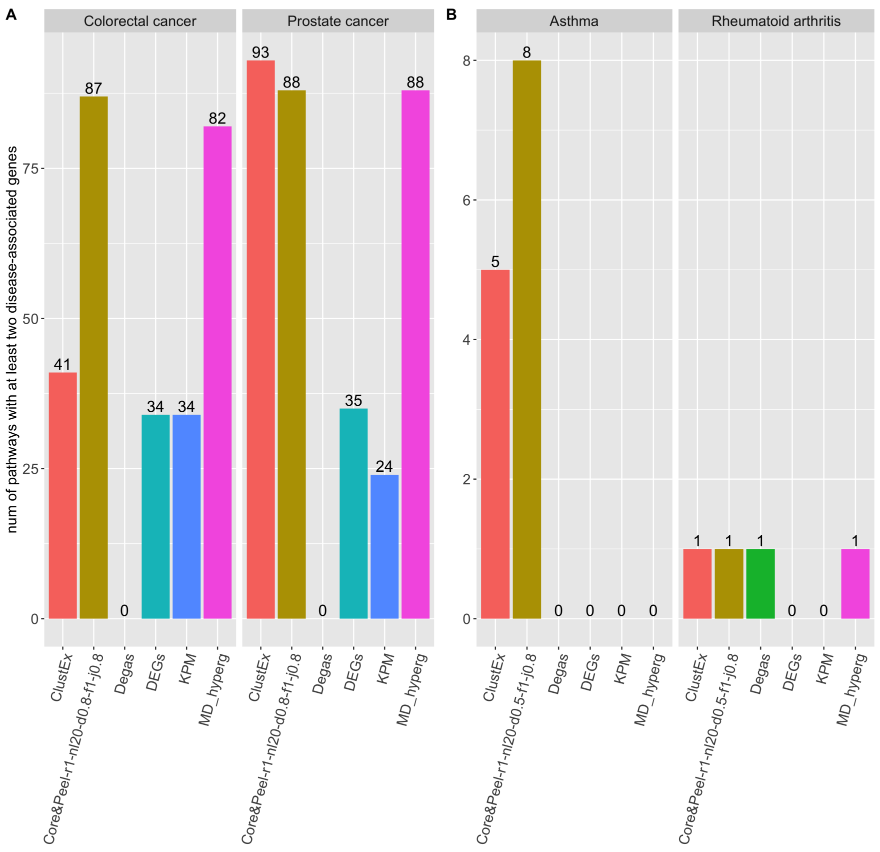

Figure 10: Number of enriched pathways with at least two disease-associated genes. The pathway enrichment analysis was performed using the Reactome database and the disease-gene association was exctracted by the DisGeNet database. (A) Prostate and Colorectal cancer cases; Core&Peel-r1-nl20-d0.8-f1-j0.8 was used. (B) Asthma and rheumatoid arthritis cases where Core&Peel-r1-nl20-d0.5-f1-j0.8 configuration was used.

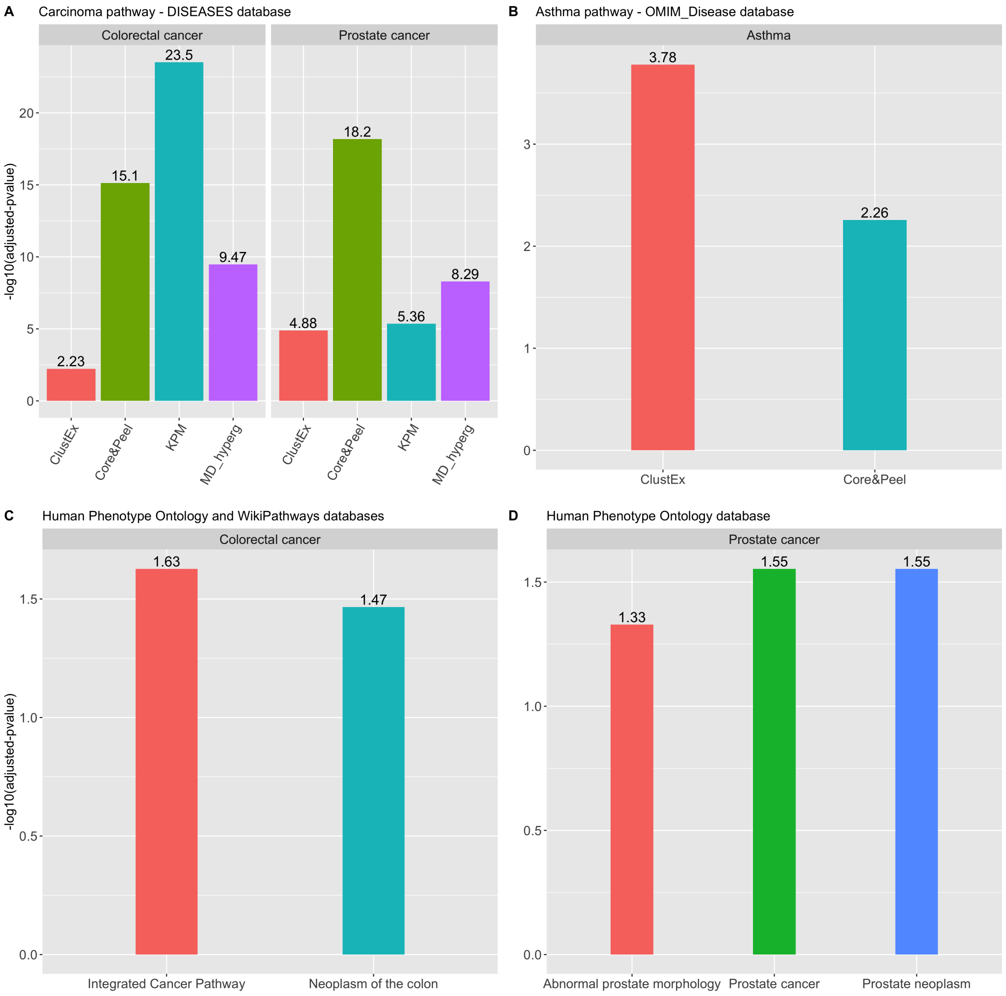

Figure 11: Results of *gProfileR* and *enrichR* analyses. The bars represent the adjusted p-value in log10-scale. (A) The adjusted p-values of Carcinoma pathway annotated in DISEASES database in colorectal and prostate cancer. Only the methods which reached the significant p-value were showed. (B) Adjusted p-values of asthma pathway annotated in OMIM_Disease database. Only Core&Peel and ClustEx modules are enriched for this pathway. (C) Adjusted p-values of Integrated Cancer Pathway and Neoplasm of colon pathways annotated in WikiPathways and Human Phenotype Ontology databases, respectively. Only Core&Peel module is enriched for these pathways in colorectal cancer. (D) Adjusted p-values of Abnormal prostate morphology, Prostate cancer and neoplasm pathways annotated in Human Phenotype Ontology database. Only Core&Peel module is enriched for these pathways in prostate cancer.

case.

| Description | p.adjust | Count | Num DisGeNet | DEGs | PMID | Reactome Category |
|---|---|---|---|---|---|---|
| Elastic fibre formation | 0.00013 | 24 | 4 | 7 | 22351925 | Extracellular matrix organization |
| Amplification of signal from the kinetochores | 0.0022 | 36 | 0 | 16 | 28651496 | Cell cycle |
| Resolution of D-loop Structuresthrough Synthesis-Dependent Strand Annealing (SDSA) | 0.0032 | 15 | 3 | 1 | 23842644 20541013 | DNA repair |
| Fatty acids | 0.0049 | 10 | 0 | 3 | 11398173 23843441 16683009 | Metabolism |
| Cell-cell junction organization | 0.0087 | 24 | 4 | 4 | 10688036 23450077 | Cell-Cell communication |
| HDR through Homologous Recombination (HRR) | 0.0088 | 25 | 3 | 2 | 30345412 24027196 | DNA repair |
| G2/M DNA replication checkpoint | 0.0123 | 5 | 0 | 2 | 25621662 18048387 | Cell cycle |
| Interleukin-36 pathway | 0.0123 | 5 | 0 | 2 | 10639195 | Immune System |
| Neurexins and neuroligins | 0.0137 | 20 | 0 | 3 | 29666812 25913192 | Neuronal System |
| ERBB2 Activates PTK6 Signaling | 0.0139 | 8 | 3 | 4 | 31294537 | Signal Transduction |
| AURKA Activation by TPX2 | 0.0249 | 27 | 1 | 4 | 22864622 30127868 28147341 | Cell cycle |
| Bile acid and bile salt metabolism | 0.0327 | 18 | 2 | 7 | 23940835 | Metabolism |
| Sodium-coupled sulphate, di- and tri-carboxylate transporters | 0.0393 | 4 | 0 | 2 | 27118869 | Transport of small molecules |
| Nuclear Receptor transcription pathway | 0.0413 | 21 | 4 | 3 | 18301781 | gene expression |

Table 10: Summary of the representative pathways of each group detected by ClueGO in prostate cancer. Description: name of the Reactome pathway; p.adjust: adjusted p-value ($< 0.05$); Count: number of genes enriched in that pathway; Num DisGeNET: number of enriched genes with a prostate-annotation in the DisGeNET database; DEGs: number of enriched genes which are differentially expressed; PMID: literature associated to that pathway; Reactome category: the Reactome category of the corresponding pathway.

#### 3.3 ClueGO analysis

In order to analyze the pathways detected only by Core&Peel in the active sub-network identification problem, we used the ClueGO [6], a Cytoscape plug-in which creates a functionally organized pathway term network. This enables us to study the most representative pathways identified by only Core&Peel. For each case study, we selected the Reactome enriched pathways detected only by Core&Peel (no overlap with any other methods) as ClueGO input. Basically a Reactome term-term similarity is calculated (for each pair of terms) using the kappa score, which takes into account the shared genes between the terms. Finally, the created network represents the terms as nodes which are linked based on a predefined kappa score level (default: 0.1). All the terms are iteratively compared and functional groups are defined. Each functional group is represented by its most significant term and the size of the nodes reflects the enrichment significance of the terms. The following networks show the Reactome-pathways identified only by Core&Peel across the four case studies and the tables summarize the representative terms of each group.

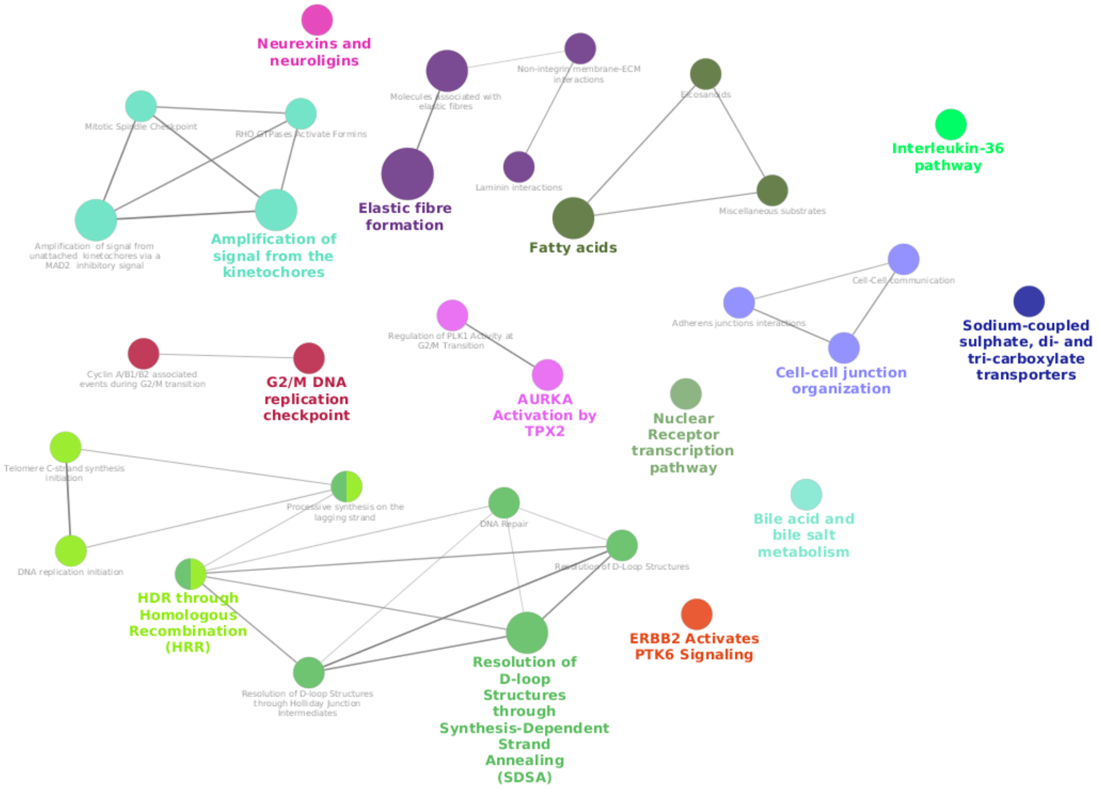

Figure 12: ClueGO results in prostate cancer case. Each functional group is highlighted by the same color and the pathways belonged to same group are linked with edges. For each group, the description of the representative pathway is colored; the descriptions of the other group members are in grey.

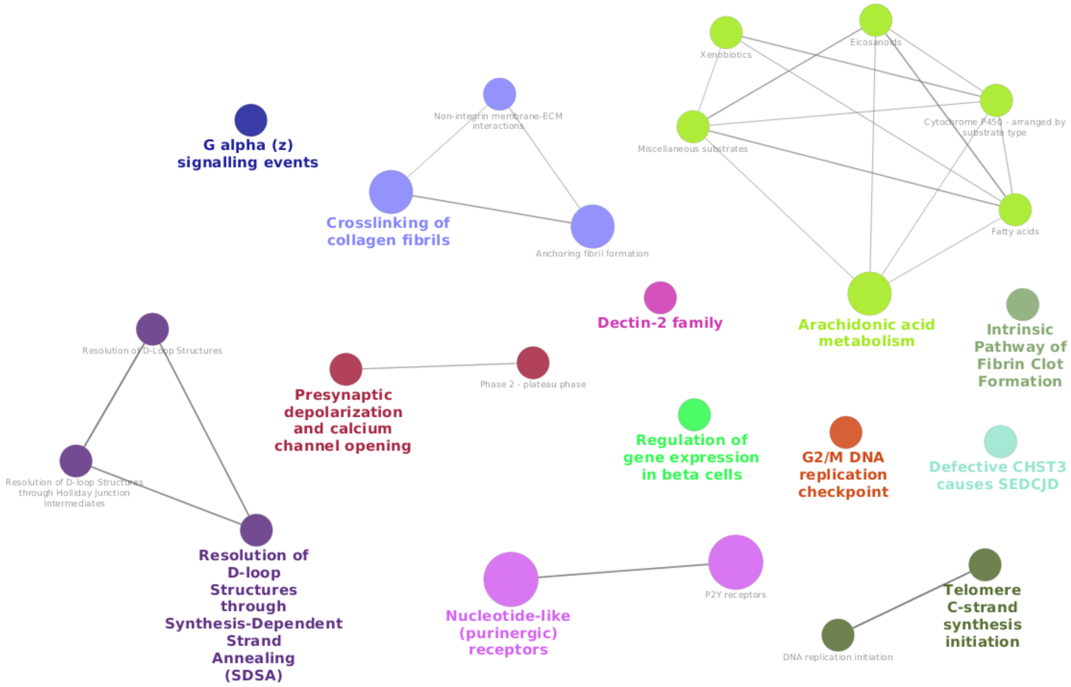

Figure 13: ClueGO results in colorectal cancer case. Each functional group is highlighted by the same color and the pathways belonged to same group are linked with edges. For each group, the description of the representative pathway is colored; the descriptions of the other group members are in grey.

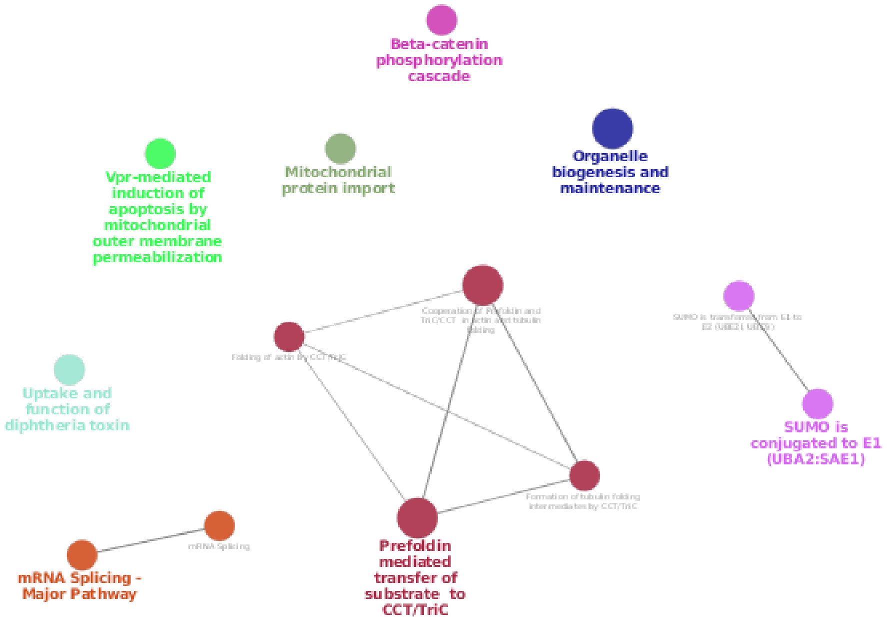

Figure 14: ClueGO results in rheumatoid arthritis case. Each functional group is highlighted by the same color and the pathways belonged to same group are linked with edges. For each group, the description of the representative pathway is colored; the descriptions of the other group members are in grey.

| Description | p.adjust | Count | Num DisGeNet | DEGs | PMID | Reactome Category |
|---|---|---|---|---|---|---|
| Nucleotide-like (purinergic) receptors | 0.0004 | 12 | 3 | 5 | 21484085 17391276 27321181 | Signal Transduction |
| Crosslinking of collagen fibrils | 0.0009 | 14 | 1 | 7 | 10972180 26940881 | Extracellular matrix Organization |
| Arachidonic acid metabolism | 0.0033 | 30 | 2 | 12 | 14619964 23715757 | Metabolism |
| Resolution of D-loop Structures through Synthesis-Dependent Strand Annealing (SDSA) | 0.0054 | 17 | 2 | 8 | 23842644 | DNA repair |
| Telomere C-strand synthesis initiation | 0.0069 | 7 | 1 | 4 | 28096526 | Cell cycle |
| Presynaptic depolarization and calcium channel opening | 0.0073 | 10 | 2 | 5 | 17111226 | Neuronal System |
| Defective CHST3 causes SEDCJD | 0.0276 | 7 | 0 | 4 | 16545347 | Disease |
| Intrinsic Pathway of Fibrin Clot Formation | 0.0313 | 14 | 0 | 4 | 23788975 8831523 | Hemostasis |
| Regulation of gene expression in beta cells | 0.0368 | 11 | 0 | 4 | | Developmental Biology |
| Dectin-2 family | 0.0368 | 11 | 0 | 5 | 26379663 30231977 | Immune System |
| G2/M DNA replication checkpoint | 0.0421 | 5 | 0 | 4 | 17241873 17943134 26805514 | Cell cycle |
| G alpha (z) signalling events | 0.0438 | 22 | 1 | 12 | 15837931 17145847 16432186 | Signal Transduction |

Table 11: Summary of the representative pathways of each group detected by ClueGO in colorectal cancer. Description: name of the Reactome pathway; p.adjust: adjusted p-value ($< 0.05$); Count: number of genes enriched in that pathway; Num DisGeNET: number of enriched genes with a prostate-annotation in the DisGeNET database; DEGs: number of enriched genes which are differentially expressed; PMID: literature associated to that pathway; Reactome category: the Reactome category of the corresponding pathway.

| Description | p.adjust | Count | num DisGeNet | DEGs | PMID | Reactome category |
|---|---|---|---|---|---|---|
| Prefoldin mediated transfer of substrate to CCT/TriC | 0.0029 | 7 | 0 | 1 | 20445014 | Metabolism of proteins |
| Organelle biogenesis and maintenance | 0.0046 | 25 | 0 | 0 | 27225300 29725325 23466882 | Organelle biogenesis And maintenance |
| Beta-catenin phosphorylation cascade | 0.0075 | 5 | 0 | 0 | 20840015 19683077 | Signal Transduction |
| SUMO is conjugated to E1 (UBA2:SAE1) | 0.0097 | 3 | 0 | 0 | 18755518 | Metabolism of proteins |
| Uptake and function of diphtheria toxin | 0.0176 | 3 | 0 | 0 | | Disease |
| mRNA Splicing - Major Pathway | 0.0247 | 17 | 0 | 0 | 26160473 21383200 | Metabolism of RNA |
| Mitochondrial protein import | 0.0248 | 8 | 0 | 0 | 15987486 | Protein localization |
| Vpr-mediated induction of apoptosis by mitochondrial outer membrane permeabilization | 0.0476 | 2 | 0 | 0 | 28102843 12813466 12094227 | Disease |

Table 12: Summary of the representative pathways of each group detected by ClueGO in rheumatoid arthritis. Description: name of the Reactome pathway; p.adjust: adjusted p-value (< 0.05); Count: number of genes enriched in that pathway; Num DisGeNET: number of enriched genes with a prostate-annotation in the DisGeNET database; DEGs: number of enriched genes which are differentially expressed; PMID: literature associated to that pathway; Reactome category: the Reactome category of the corresponding pathway.

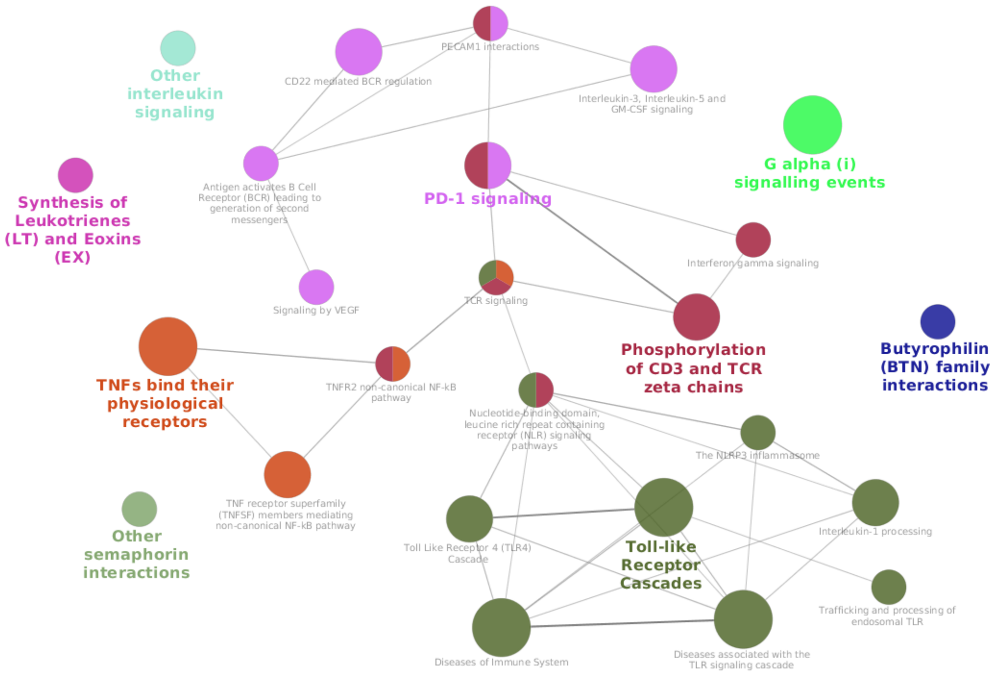

Figure 15: ClueGO results in asthma case. Each functional group is highlighted by the same color and the pathways belonged to same group are linked with edges. For each group, the description of the representative pathway is colored; the descriptions of the other group members are in grey.

| Description | p.adjust | Count | num DisGeNet | DEGs | PMID | Reactome category |
|---|---|---|---|---|---|---|
| TNFs bind their physiological receptors | 1.48e-05 | 14 | 0 | 1 | 18036647 17475560 | Immune System |
| Toll-like Receptor Cascades | 0.00011 | 37 | 1 | 2 | 27543676 28692144 | Immune System |
| G alpha (i) signalling events | 0.00016 | 66 | 4 | 5 | 18278482 18552456 | Signal Transduction |
| Phosphorylation of CD3 and TCR zeta chains | 0.00061 | 10 | 1 | 0 | 99221231 0375551 | Immune System |
| PD-1 signaling | 0.00099 | 10 | 1 | 0 | 18173375 28865147 30644695 | Immune System |
| Other interleukin signaling | 0.00597 | 10 | 0 | 0 | 26750312 15464449 23283176 | Immune System |
| Butyrophilin (BTN) family interactions | 0.00686 | 6 | 0 | 0 | 22030238 20944003 | Immune System |
| Other semaphorin interactions | 0.01838 | 8 | 0 | 0 | 23994348 23932461 | Developmental Biology |
| Synthesis of Leukotrienes (LT) and Eoxins (EX) | 0.02707 | 8 | 2 | 0 | 20920774 21059119 | Metabolism |

Table 13: Summary of the representative pathways of each group detected by ClueGO in asthma. Description: name of the Reactome pathway; p.adjust: adjusted p-value ($< 0.05$); Count: number of genes enriched in that pathway; Num DisGeNET: number of enriched genes with a prostate-annotation in the DisGeNET database; DEGs: number of enriched genes which are differentially expressed; PMID: literature associated to that pathway; Reactome category: the Reactome category of the corresponding pathway.
